# AI-Enforced Ultra-Large Virtual Screening Discovers Potent CD28 Binders

**DOI:** 10.64898/2026.03.26.714621

**Authors:** Saurabh Upadhyaya, Michele Roggia, Shaoren Yuan, Sandro Cosconati, Moustafa T. Gabr

## Abstract

Targeting protein-protein interactions (PPIs) with small molecules is historically challenging due to shallow, solvent-exposed interfaces that lack classical binding pockets. Furthermore, employing traditional structure-based virtual screening (SBVS) across ultra-large chemical spaces to find novel chemotypes imposes prohibitive computational bottlenecks. Here, we report the first prospective, real-world application of the PyRMD2Dock platform, an AI-enforced SBVS workflow that integrates machine learning and standard docking available within the PyRMD Studio suite. To target the structurally demanding immune receptor CD28, a chemically diverse subset of 2.4 million molecules from the Enamine REAL Diversity Space was docked into a cleft adjacent to the canonical ligand interface. These data were used to train 672 classification models, and the optimized model rapidly screened the remaining ∼46 million compounds. Following interaction filtering and clustering, 232 highly prioritized ligands were identified. Experimental validation of 150 purchased candidates yielded a remarkable hit rate, identifying multiple direct CD28 binders. Lead compounds **100** and **104** exhibited submicromolar affinity (K_d_ = 343.8 nM and 407.1 nM, respectively), potent CD28-CD80 disruption, and functional blockade in cellular reporter assays. Furthermore, these compounds successfully reduced cytokine secretion in primary human tumor-PBMC and epithelial tissue co-culture models. This study validates PyRMD2Dock as a highly scalable, effective protocol for mining massive chemical libraries to discover small-molecule modulators of challenging immune receptor interfaces.

## Introduction

Targeting protein–protein interactions (PPIs) using small molecules remains a formidable challenge in computational drug discovery due to the prevalence of broad, solvent-exposed, and topographically shallow binding interfaces.^1, 2^ Unlike enzymes or receptors that typically feature well-defined, deep orthosteric cavities, PPI surfaces frequently lack deep concave pockets suitable for classical structure-based ligand design.^3, 4^

The costimulatory immune receptor CD28 serves as a prime example of such a structurally demanding PPI target. CD28 is a homodimeric immunoglobulin-like receptor that engages the ligands CD80 and CD86 through a largely planar and solvent-accessible extracellular β-sandwich surface.^5, 6^ The absence of a deeply recessed cavity in this canonical interface makes CD28 an instructive model system for evaluating computational strategies aimed at exploiting shallow surface depressions and partially lipophilic microenvironments.^7^ These architectural constraints have historically limited the development of small-molecule modulators and position CD28 as a structurally informative model for structure-guided targeting of flat immune receptor interfaces.^8,9^

To productively target such intractable interfaces with small molecules, systematic structural interrogation is essential to identify transient or subtle ligandable microenvironments.^10^ Concurrent with these structural analyses, recent computational advances emphasize that screening ultra-large molecular databases significantly increases the probability of discovering novel chemotypes for challenging targets.^11^ However, deploying traditional structure-based virtual screening (SBVS) workflows across billions of compounds imposes prohibitive computational bottlenecks. To overcome these scaling limitations, we utilized PyRMD2Dock within the PyRMD Studio suite.^12^ This comprehensive graphical user interface (GUI) democratizes sophisticated AI workflows, allowing researchers to perform both ligand-based and structure-based virtual screening without requiring advanced coding expertise. PyRMD2Dock bridges the gap between AI-driven ligand-based virtual screening (LBVS) and SBVS by coupling the machine learning classifier PyRMD with the high-performance docking engine AutoDock-GPU (AD4-GPU).^12,13^ Furthermore, PyRMD Studio incorporates critical code optimizations that increase screening speed by more than 3-fold compared with earlier versions.

By training its predictive machine learning model on the docking scores of a small, randomly sampled subset of the chemical library, PyRMD learns to identify chemical features correlated with favorable binding. This methodology enables rapid, effective screening of ultra-large chemical spaces without the immense computational power required to dock every compound explicitly. In this study, we showcase the specific utility of the PyRMD2Dock platform in identifying potent small-molecule disruptors of the structurally challenging CD28–B7 interaction. To experimentally assess structure-guided ligand prioritization, computationally selected CD28 ligands were evaluated using a tiered validation framework that integrated quantitative biophysical measurements, interface-disruption assays, and functional signaling analyses. Direct molecular engagement was first examined using microscale thermophoresis and complementary time-resolved interaction assays to determine equilibrium dissociation constants and confirm saturable binding behavior. Functional perturbation of the CD28–B7 interface was subsequently evaluated using competitive recombinant binding assays and protein–protein complementation systems, enabling quantitative assessment of interface disruption.

Compounds demonstrating reproducible binding and competitive inhibition were further interrogated in CD28-dependent signaling models to establish downstream biological consequences. Reporter-based blockade bioassays quantified suppression of costimulatory signaling, while human immune cell–based systems—including tumor–PBMC and mucosal epithelial–PBMC co-culture platforms—enabled evaluation of cytokine modulation under physiologically relevant activation conditions. By linking our AI-enforced virtual screening cascade directly to measurable immunomodulation, these findings not only validate the PyRMD2Dock protocol but also demonstrate that shallow, solvent-exposed immune receptor interfaces can be successfully targeted by small molecules, thus broadening the tractable landscape of complex PPIs.

## Results and Discussion

### AI-Enforced Structure-Based VS against CD28

To discover novel CD28 binders, the PyRMD Studio suite, through its PyRMD2Dock^12^ protocol, was employed as shown in Figure 1.

**Figure 1.**
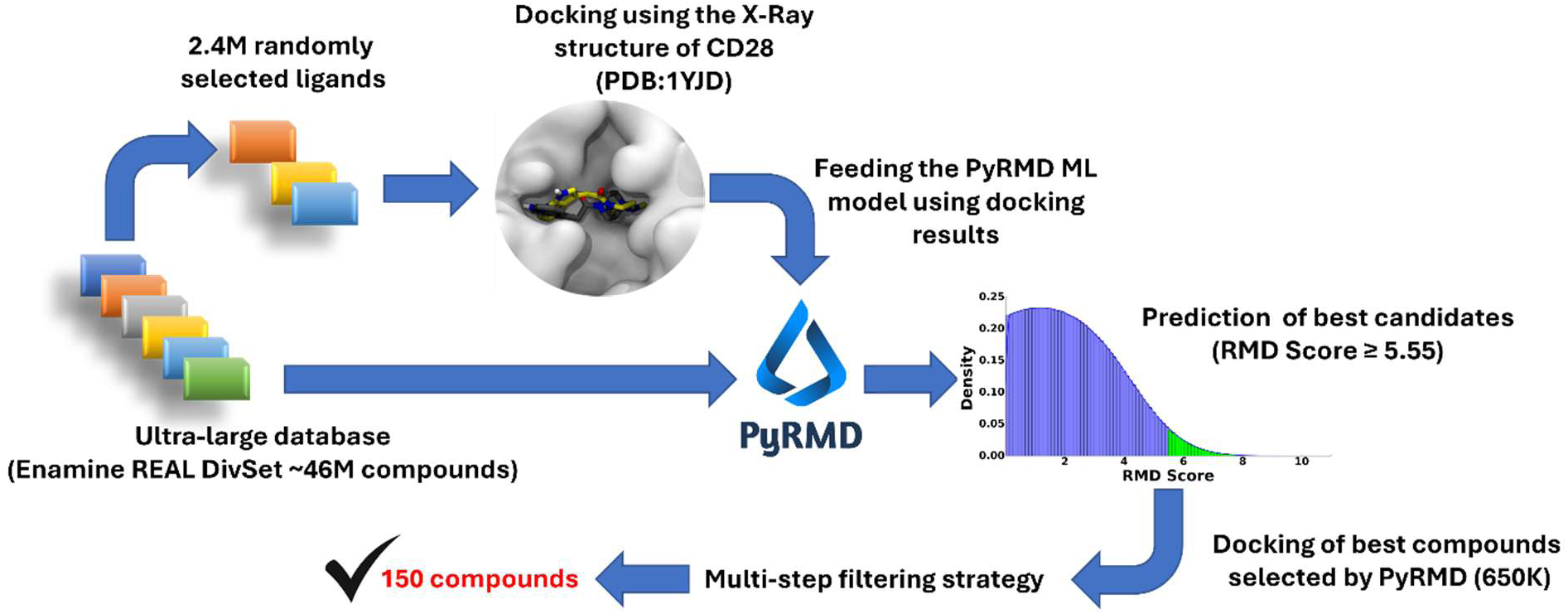
The adopted PyRMD2Dock workflow.

It combines the Ligand-Based VS (LBVS) tool PyRMD^13^ with one of the widely used docking software, AutoDock-GPU (AD4-GPU).^12, 13^ By using the PyRMD2Dock pipeline, we established a robust workflow capable of ultra-fast screening across massive chemical libraries to pinpoint molecules with the highest predicted binding affinity to a target protein. The initial phase of this protocol involved the preparation and molecular docking of a subset of 2.4 million compounds for training purposes from the Enamine REAL DivSet database. To ensure a highly representative and chemically diverse dataset, these compounds were selected using an RDKit-based diversity picking approach. Specifically, molecular 2D structures were generated from SMILES strings, and Extended Connectivity Fingerprints with a radius of 3 (ECFP6) were computed. A MaxMin algorithm,^15^ relying on Tanimoto distances, was then applied to extract the optimal, maximally diverse subset. This subset was then docked into the X-Ray structure of human CD28 in complex with the Fab fragment of a mitogenic antibody (PDB ID: 1YJD).^6^ Here, the primary ligand-binding site of this glycoprotein is defined by the MYPPPY motif (Figure 2a), spanning amino acids 99-104. However, a close inspection of the CD28 3D structure revealed that this canonical binding region is relatively flat and lacks a cavity deep enough to effectively accommodate and stably bind small-molecule drugs. Thus, our structural analysis was redirected towards a distinct pocket adjacent to the natural ligand-binding region. This cavity is delimited by a specific set of amino acids, namely H38, F93, K95, D106, N107, K109, and S110 (Figure 2a-b).

**Figure 2.**
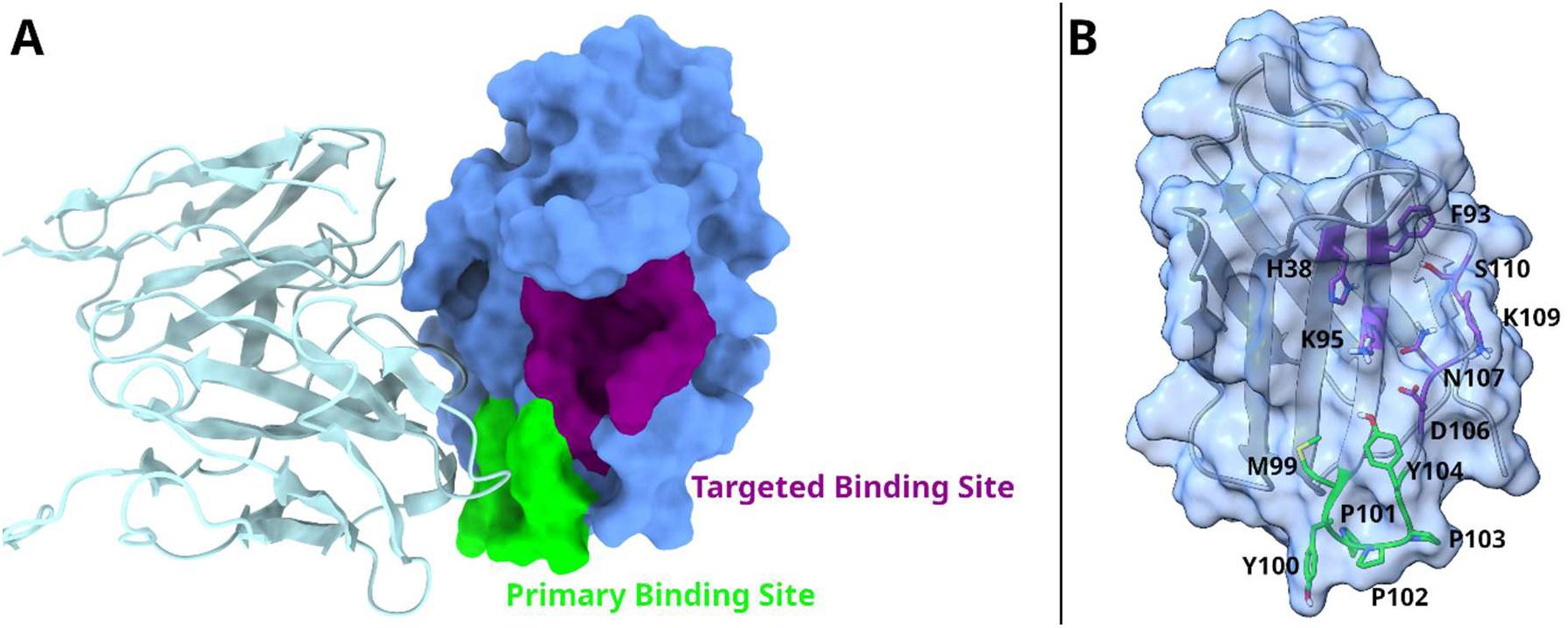
A) Overall structure of CD28 (blue surface) in complex with the Fab fragment (sky blue ribbons) where the primary binding site (green), and the targeted cleft (violet) are highlighted. B) A detailed, semi-transparent view of the CD28 monomer revealing the specific amino acid residues involved in these regions.

To evaluate the structural relevance of this pocket, we investigated the features of CTLA-4, a receptor sharing high structural homology with CD28. Specifically, we analyzed the 3D structure of CTLA-4 in complex with the Fab fragment of the monoclonal antibody tremelimumab (PDB ID: 5GGV).^16^ Structural data indicate that the HCDR3 loop of tremelimumab (residues 101–110) is heavily involved in binding CTLA-4. Notably, this specific portion of the antibody interacts with a CTLA-4 region that corresponds to the putative cleft we elected as the target region in CD28. Based on these structural alignments and the established interaction of tremelimumab with the adjacent cavity on CTLA-4, the corresponding pocket identified on CD28 was selected as a viable target for our virtual screening campaign.

Following the docking of the training set, for all the compounds the predicted lowest binding free energy (ΔG_AD4_) was extracted, and the results were analyzed based on their ΔG_AD4_ values distribution. Here, the compounds were separated as “active” and “inactive” depending on their docking score. Specifically, four distinct activity thresholds for the active compounds were established: −8.25, −8.0, −7.75, and −7.50 kcal/mol. Any compound scoring at or below these limits was assigned to the “active” group. Conversely, three thresholds were selected for the inactive compounds: −4.75, −4.50, and −4.25. Molecules with a predicted ΔG_AD4_ at or above these cutoffs were classified as “inactive”. By using these parameters, 12 possible dataset combinations (4 active thresholds × 3 inactive thresholds) were generated to conduct a thorough benchmarking analysis.

A critical step for the success of the entire workflow is the optimization of the training parameters, particularly the ε cutoff values for both “active” and “inactive” classes. These cutoffs define the maximum allowed projection distance for an unclassified compound within the model’s linear subspace, thus heavily influencing the final classification accuracy. To identify the optimal settings, the ε cutoff values ranging from 0.99 down to 0.70 (with a step of 0.05) for the active sets, and from 0.95 down to 0.60 (with a step of 0.05) for the inactive sets were systematically evaluated (i.e., 672 different models were evaluated). Here, the employment of the PyRMD Studio suite was instrumental, as it implements a cross-validation adapted version of the StratifiedGroupKFold splitting utility, coupled with Butina clustering. This robust validation strategy ensures that no examples from the same structural cluster are present in both the training and test sets, thereby effectively mitigating data leakage and preventing overly optimistic performance estimates. Thus, we evaluated the performance of all generated models by analyzing a combination of key classification metrics. While maximizing the TPR/FPR tradeoff was a primary criterion, it was equally crucial to ensure high Precision and a strong F-score. Therefore, we designed our selection strategy to identify the configuration that offered the best overall balance, prioritizing the model capable of providing an excellent TPR/FPR trade-off alongside robust Precision and F-score metrics. This balance was achieved by the model defined by an “active” threshold of −7.75 kcal/mol, an “inactive” threshold of −4.75 kcal/mol, and ε cutoffs of 0.95 and 0.70 for “actives” and “inactives”, respectively (Supporting Information file “benchmark_results_all.xlsx”). Consequently, this specific setup was selected for the large-scale screening phase.

Using the optimized PyRMD model trained on these docking results, the Enamine REAL Diversity Set (ERDS), a large-scale database comprising 48 million compounds, was screened. The PyRMD2Dock screening module of PyRMD Studio successfully processed this library, ultimately identifying 20,163,675 compounds classified as potential CD28 binders. Notably, refactoring critical internal operations to vectorized equivalents in PyRMD Studio enabled this massive screening to be completed with an approximately 3.3-fold increase in processing speed relative to the previous software version. These results were somewhat expected, considering that the training data for the PyRMD model were derived from docking calculations. In fact, docking simulations rely on approximations and scoring functions that can yield a significant number of false positives, thereby introducing noise. Thus, we applied a strict cut-off to this massive output, thereby selecting only the top-ranking 650,000 compounds from this extensive screening, exhibiting RMD^12,13^.values spanning from 10.98 to 5.55.

To validate the predictive capability of the model, the distribution of the ΔG_AD4_ values of the selected compounds was compared to that obtained by docking an equal number of compounds randomly chosen from the database (Figure 3a). This analysis revealed a clear enrichment in low ΔG_AD4_-scores within the PyRMD selection. As expected, an overlap between the two distribution curves is present, occurring in proximity to the −7.00 kcal/mol value. This is due to the intrinsic error connected with the docking scoring function, which affects the training process. Interestingly, the mean and the mode of the ΔG_AD4_ values obtained when docking the PyRMD-selected ligands (−7.48 and −7.43 kcal/mol, respectively) well approximate the active cutoff ΔG_AD4_ value employed in the training process (−7.75 kcal/mol). This outcome is again expected as during the learning training process, the compounds defined as "actives" provided the model with the specific chemical features responsible that in docking calculations contribute to a ΔG_AD4_ ≤ −7.75 kcal/mol. Consequently, PyRMD naturally selects new compounds possessing those same structural determinants, resulting in a large population of molecules that achieve similar docking scores while still successfully enriching the overall selection with better-scoring compounds.

**Figure 3.**
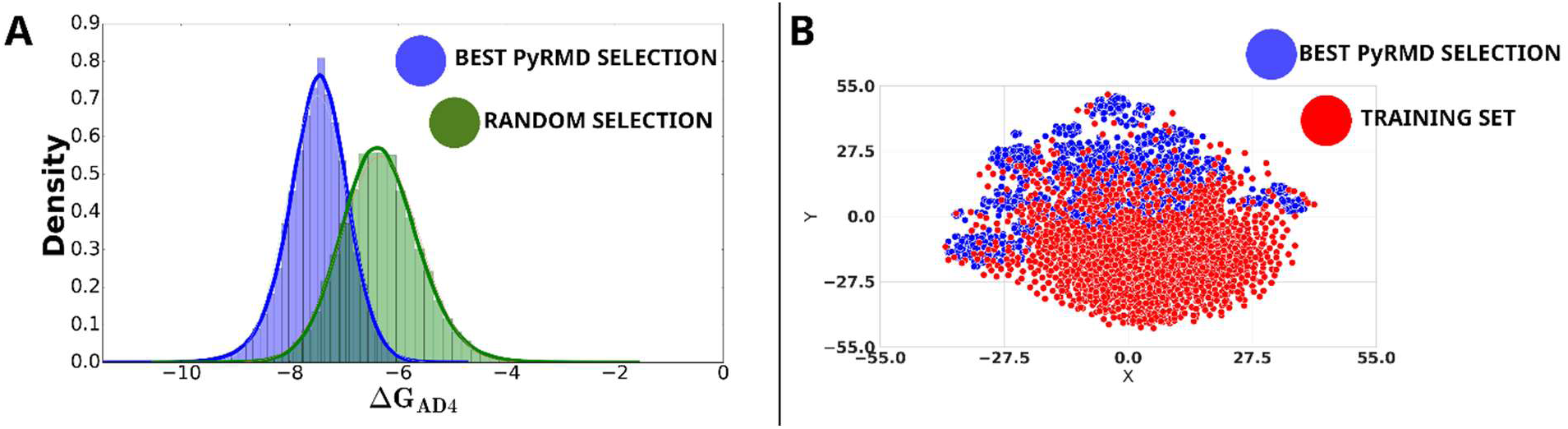
A) Distribution of the Δ*G*_AD4_ values obtained when docking the best-scoring compounds from PyRMD selection (blue) and the same number of compounds randomly selected from the same dataset (green). B) Chemical space of the PyRMD training and output subsets. The training active compounds are shown in red (labeled as “TRAINING SET” in the legend), while the subset consisting of the top screened compounds with the highest RMD scores is shown in blue (labeled as “BEST PyRMD SELECTION” in the legend).

Although this analysis provides a clear indication that the RMD can be effectively trained using docking results to predict the outcome of a virtual screening campaign, it does not provide insight into the coverage of chemical space in the output compared to the input. To address this, an analysis was conducted on the chemical space covered by a representative subset of the training set and an equal number of compounds with the highest RMD score output, using t-distributed stochastic neighbor embedding (t-SNE) plots (Figure 3b). The t-SNE analysis highlighted that the training dataset derived from docking is intrinsically noisy, as indicated by the utter absence of chemical clusters of compounds. However, the ML algorithm, by implementing the Marchenko-Pastur distribution, effectively denoises the docking data and facilitates the extraction of meaningful signals. Consequently, PyRMD Studio was able to successfully enrich discrete portions of the training set, identifying families of compounds featuring similar chemical features and disregarding chemical information present in the training set that is identified as noise.

Thus, the best 650,000 ligands returned by PyRMD underwent docking calculations into the same CD28 structure employed for the training set.^13^ Based on the predicted ΔG_AD4_ values, only compounds presenting an energy score ≤ −7.75 kcal/mol were advanced to the next stage, reducing the pool to ∼200,000 molecules. To further refine this selection and guarantee that the remaining compounds correctly fit within the identified cleft, a structure-based filter approach was applied. We defined a filtering criterion based on two strategic interaction points spanning the targeted cavity: K109 and C94. The first one was selected as it is one of the key residues delimiting the pocket (Figure 2) and its positively charged side chain should serve as an excellent attachment point, able to establish strong ionic interactions with the ligand candidates. On the other hand, although C94 does not delimit the targeted binding site, it is centrally located at the deep bottom of the pocket.^6^ Thus, filtering out all molecules that failed to establish simultaneous contacts with both residues ensured the selection of ligands strictly occupying the full volume of the targeted cavity. This interaction-based filtering step yielded ∼94,000 ligands, which were subsequently subjected to a solvent-accessible surface area (SASA) analysis using Maestro (Schrödinger) as well as the ΔSASA to quantify the steric fit of the docked compounds. This latter metric represents the ligand’s buried surface area upon binding, calculated as the difference between the SASA of the free ligand and its SASA within the receptor-ligand complex. To narrow down the selection while maintaining chemical diversity, the molecules were clustered based on their structural features. From the various generated clusters, we selected multiple compounds rather than strictly limiting the choice to the single top-scoring molecule per cluster. This selection process was primarily driven by the ΔSASA values, allowing us to come up with a subset of ∼13,000 diverse ligands that exhibited the optimal fit within the pocket. At this stage, visual inspection of the remaining ∼13,000 ligands revealed the presence of three distinct binding orientations within the targeted (Figure 4).

**Figure 4.**
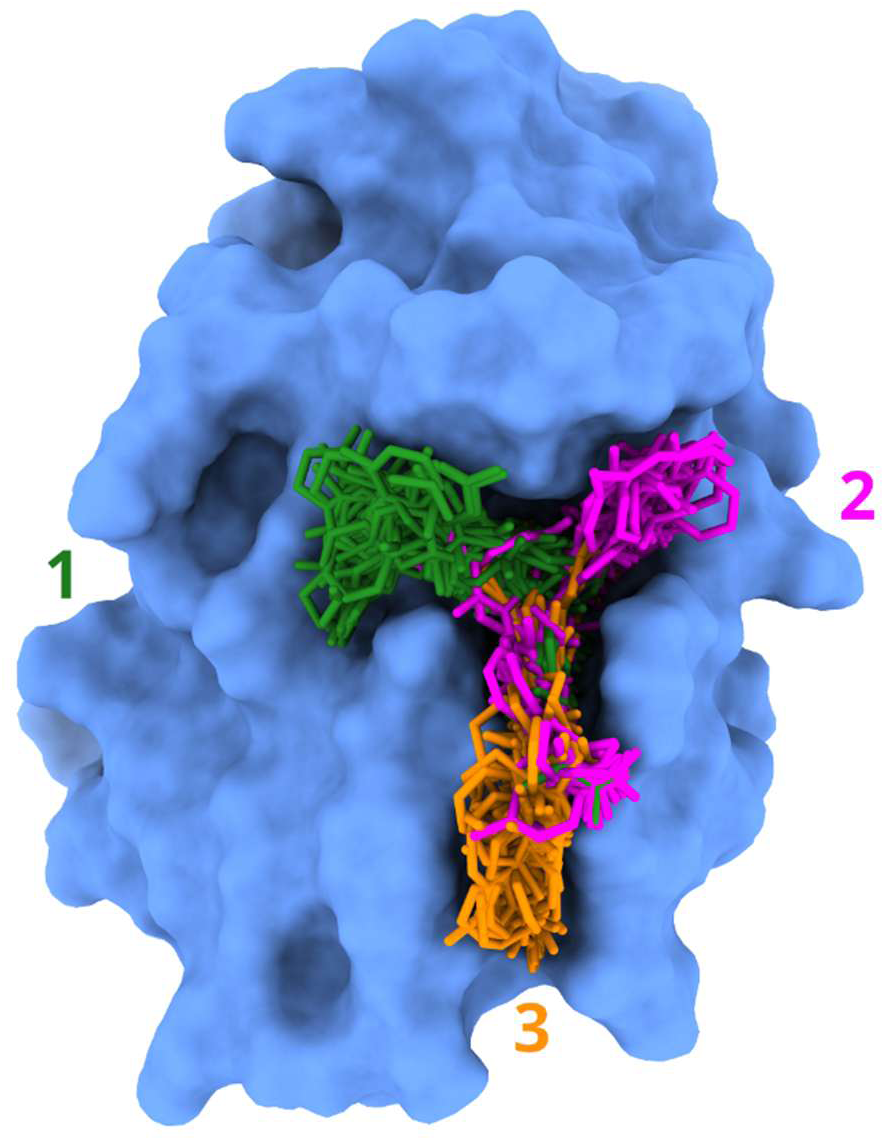
Representative subset of ligands illustrating the three distinct binding orientations identified within the targeted binding site from a pool of ∼13,000 compounds. The compounds belonging to the different binding modes are represented as green, magenta, and orange sticks for clusters 1, 2, and 3, respectively, while the protein is depicted as a blue surface.

Subsequently, to ensure a comprehensive evaluation of these orientations, iterative clustering rounds were performed separately for each of the three groups, categorized by binding mode. Ultimately, this approach led to a final, highly refined selection of 232 ligands. Based on these computational results, 150 compounds were successfully purchased and subjected to biological assays.

### Primary Dianthus Screening of 150 Computationally Selected Compounds

Following large-scale virtual screening of approximately 48 million compounds (described above), a prioritized subset of 150 chemically diverse molecules was selected for experimental evaluation. These compounds were assessed using a Dianthus-based assay employing the purified extracellular domain (ECD) of CD28 to detect compound-induced perturbations in the assay signal under defined biochemical conditions.

Vehicle control wells (2% DMSO in PBST, 0.05% Tween) established a stable baseline signal with a mean Fnorm of 0.863. The reference compound BPU11 reduced the signal to a mean Fnorm of 0.769, confirming that small-molecule engagement of CD28 ECD produces a measurable change within the assay window.^17^

To identify statistically significant signal-altering events, a stringent ±5 standard deviation threshold relative to the vehicle mean was applied, corresponding to an Fnorm interval of 0.828–0.894. The majority of screened compounds generated Fnorm values within this confidence range, indicating no detectable interaction under the assay conditions.

In contrast, twelve compounds reproducibly produced Fnorm values outside the predefined statistical interval. These molecules were therefore classified as primary hits, corresponding to approximately 7.3% of the tested subset (Figure 5A). Given that the assay contains only purified CD28 ECD and small molecules, these signal deviations reflect compound–protein interactions sufficient to alter the Dianthus readout.

The twelve primary hits were subsequently advanced to orthogonal biophysical validation by microscale thermophoresis (MST) to quantify direct binding affinity and assess concentration-dependent engagement of CD28.

**Figure 5.**
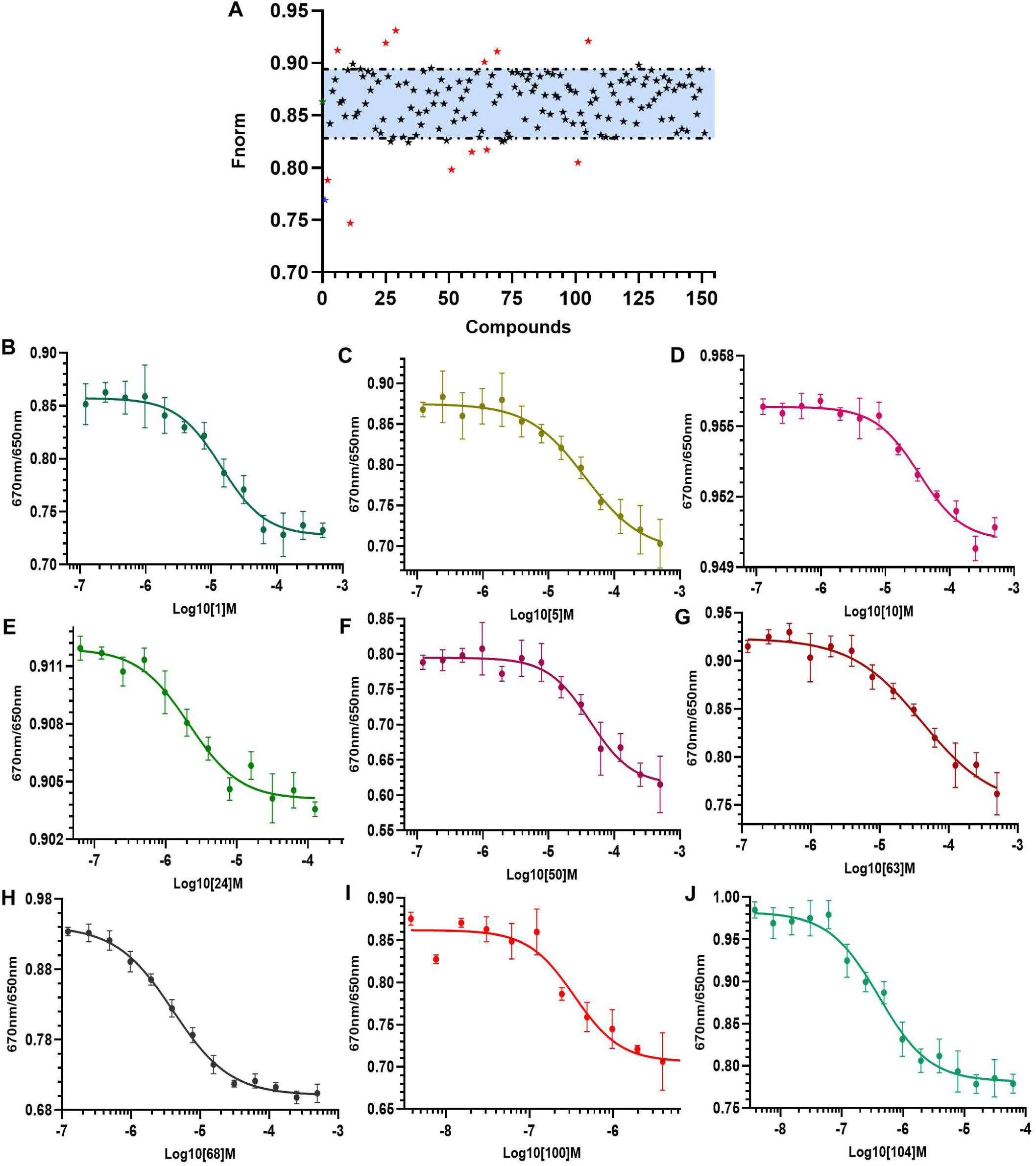
(A) Dianthus-based screening of 150 computationally prioritized compounds using purified CD28 extracellular domain (ECD). Individual compounds are plotted as Fnorm values. The shaded region represents the ±5 SD interval relative to the vehicle control (2% DMSO in PBST, 0.05% Tween). Compounds outside this interval were classified as primary hits. BPU11 is shown as the positive control. (B–J) Concentration–response binding curves obtained by microscale thermophoresis (MST) using purified CD28 ECD. Compounds were tested over a concentration range of 500 μM to 1 nM (1:1 serial dilution) with 2% DMSO maintained across all conditions. Data are shown as mean ± SEM. Solid lines represent nonlinear regression fits used to calculate equilibrium dissociation constants (K_d_). (B) **1**, (C) **5**, (D) **10**, (E) **24**, (F) **50**, (G) **63**, (H) **68**, (I) **100**, and (J) **104.**

### Orthogonal Validation of CD28 Binding by MST

To determine whether Dianthus-derived hits reflected direct engagement of CD28, binding affinities were quantified using microscale thermophoresis (MST) with purified CD28 extracellular domain (ECD). Compounds were evaluated over a concentration range spanning 500 μM to 1 nM in a 1:1 serial dilution format, with 2% DMSO maintained across all conditions. Binding measurements were performed in three independent experiments to ensure reproducibility.

Concentration-dependent thermophoretic shifts yielded well-defined sigmoidal binding curves for multiple compounds. Two candidates, **100** and **104**, exhibited submicromolar affinity with dissociation constants (K_d_) of 343.78 ± 22.6 nM and 407.08 ± 14.9 nM, respectively. **24** and **68** demonstrated low-micromolar binding (K_d_ = 2.21 ± 1.4 μM and 4.05 ± 3.2 μM), while 1 bound with moderate affinity (K_d_ = 14.16 ± 3.2 μM). Several additional compounds displayed mid-micromolar engagement (**5**, **10**, **50**, **63**, **64**, **58**; K_d_ ≈ 36–85 μM), whereas **28** showed substantially weaker binding (K_d_ = 426.9 ± 29.2 μM) (Figure 4B-J and Figure S19).

This orthogonal validation step refined the initial Dianthus hit set into a ranked affinity series, identifying **100** and **104** as the highest-affinity CD28 binders for subsequent functional and mechanistic evaluation.

### Competitive Disruption of CD28–CD80 Interaction by Compounds

To evaluate whether MST-confirmed CD28 binders disrupt ligand recognition, all eleven compounds demonstrating dose-dependent binding in MST were assessed in a biochemical CD28–CD80 competition ELISA. Recombinant CD28 was immobilized on 96-well plates and incubated with biotinylated CD80 in the presence of increasing compound concentrations. Inhibition curves were generated from three independent experiments to derive IC₅₀ values.

Seven of the eleven MST-positive compounds exhibited reproducible, concentration-dependent inhibition of CD28–CD80 binding. **100** showed the highest potency (IC₅₀ = 231.4 ± 34 nM), followed by **104** (IC₅₀ = 840.6 ± 112.8 nM) and **68** (IC₅₀ = 1.32 ± 0.8 μM). **1** and **24** displayed moderate inhibition (IC₅₀ = 9.5 ± 1.6 μM and 22.3 ± 6.1 μM), whereas **10** showed weaker activity (IC₅₀ = 105.8 ± 12.65 μM) (Figure 6A-F) and **28** exhibited minimal inhibition (IC₅₀ = 411.3 ± 51.7 μM) (Figure S20 in Supporting Information).

**Figure 6.**
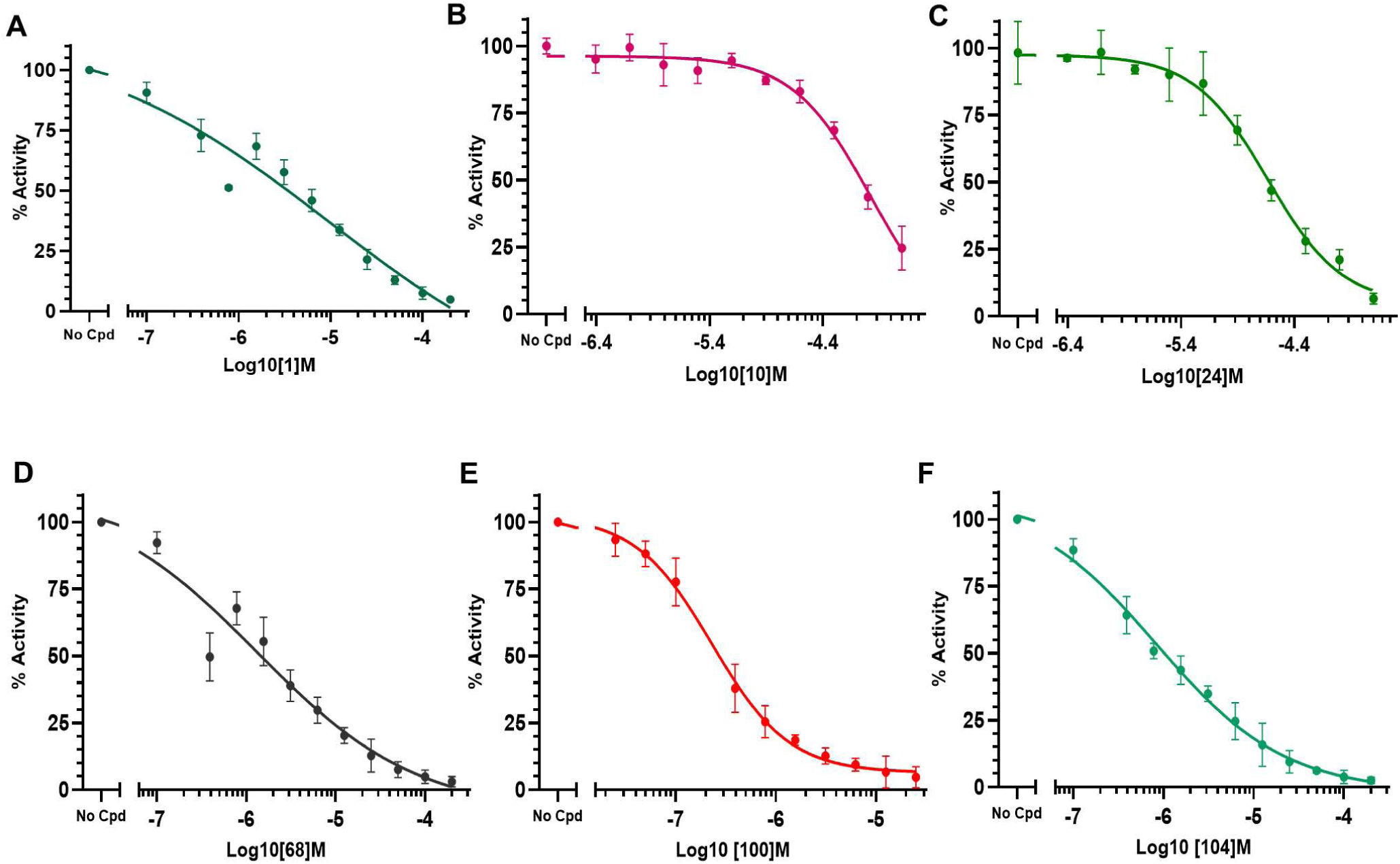
Dose–response curves from ELISA-based CD28–CD80 binding assays. CD28 extracellular domain was immobilized and incubated with biotinylated CD80 in the presence of increasing concentrations of the indicated compounds. Percent activity was calculated relative to the no-compound control (set to 100%). Compounds were tested over a defined concentration range using serial dilutions. Data are presented as mean ± SEM. Solid lines represent nonlinear regression fits used to determine IC₅₀ values. (A) **1**, (B) **10**, (C) **24**, (D) **68**, (E) **100**, and (F) **104**.

The remaining MST-confirmed binders did not demonstrate reproducible dose-dependent blockade under identical conditions, indicating that CD28 binding does not uniformly translate into competitive disruption of CD80 engagement. Notably, the most potent functional inhibitors (**100** and **104**) were also the highest-affinity binders identified by MST, revealing a clear correlation between binding strength and competitive activity. Table 1 summarizes the results of the biological evaluation of the positive hits.

**Table 1.**
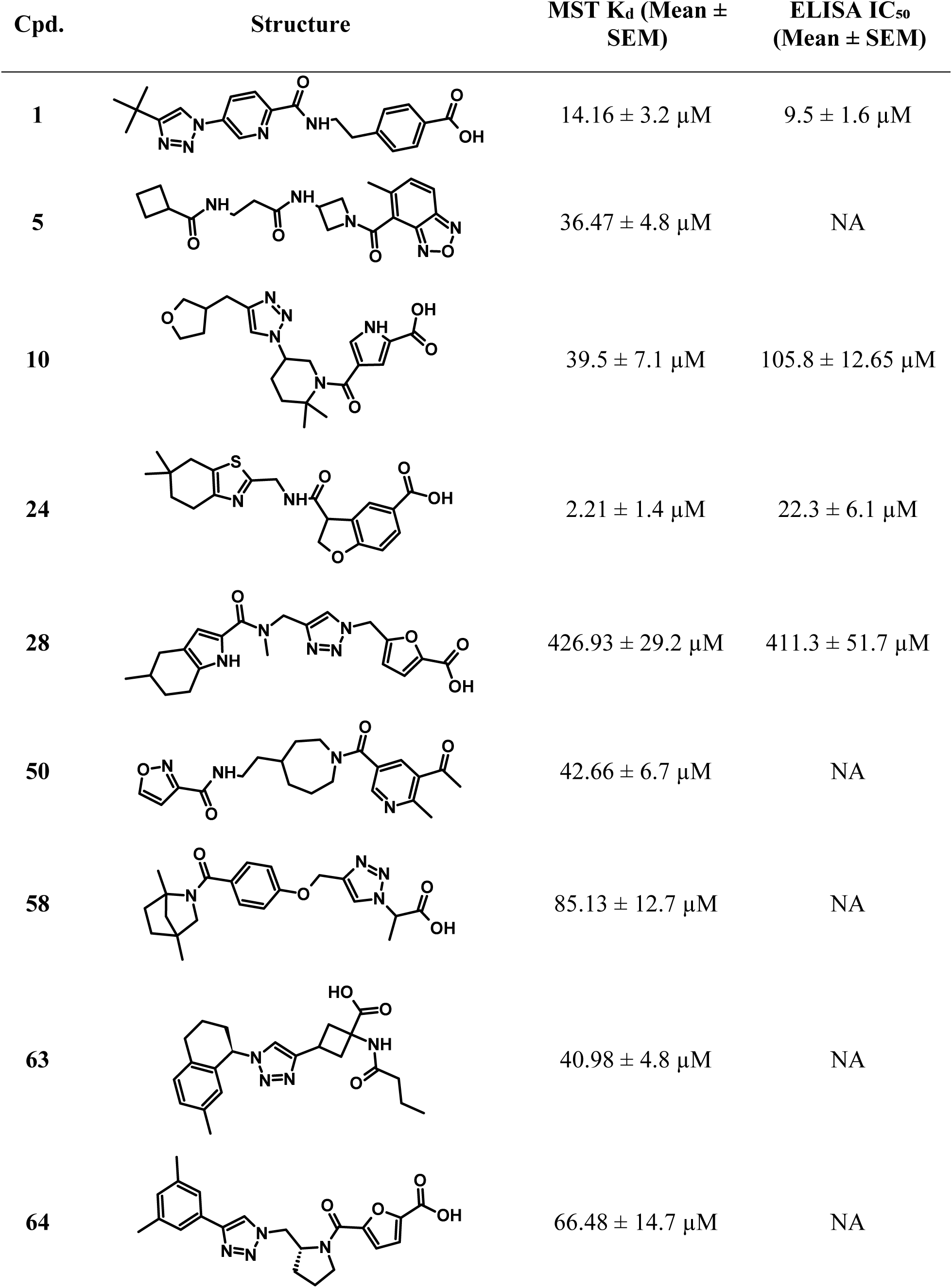
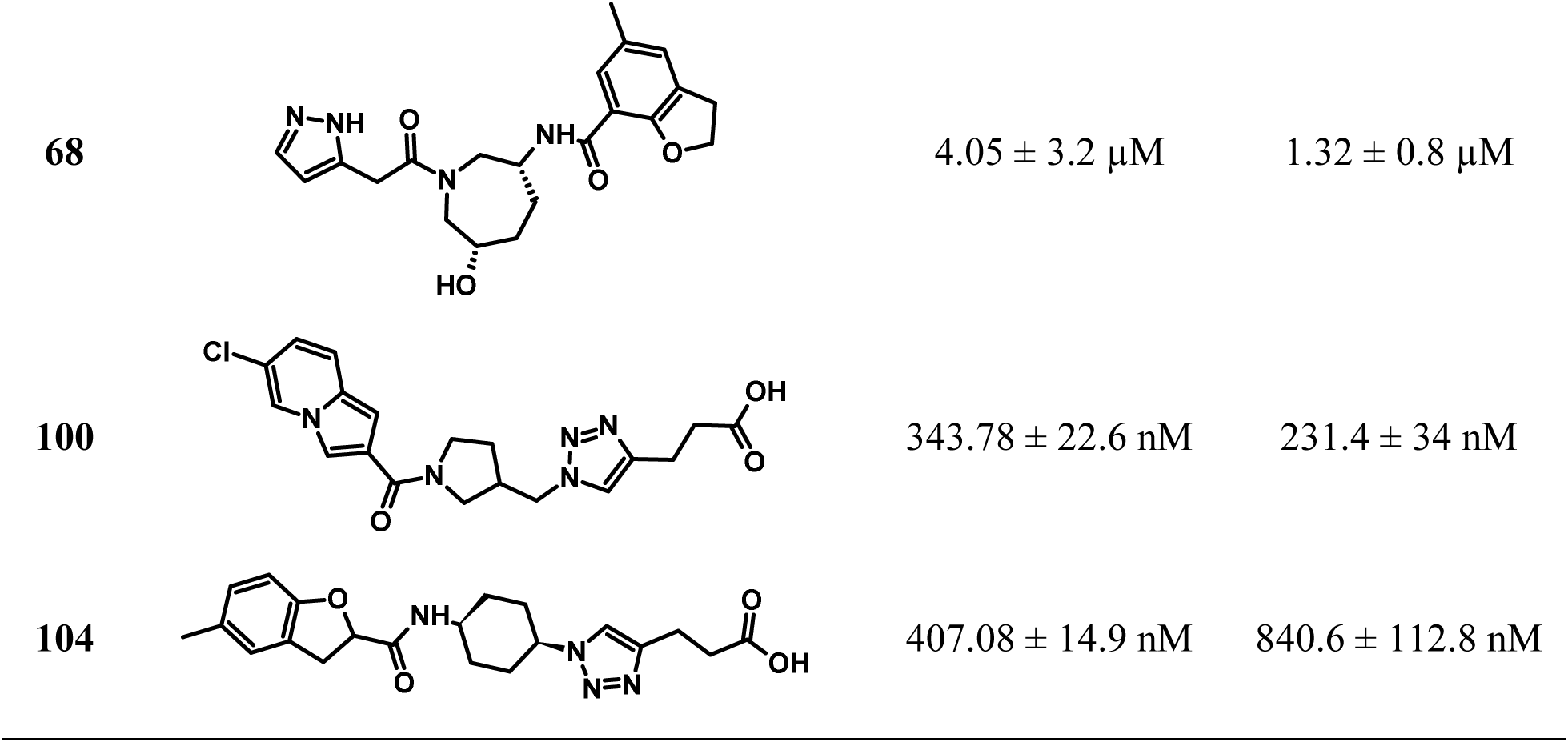
Structures of the best compounds from among the 150 tested ones, along with their MST and ELISA K_d_, and IC_50_ values, respectively.

| Cpd. | Structure | MST K<sub>d</sub> (Mean ± SEM) | ELISA IC<sub>50</sub> (Mean ± SEM) |
| --- | --- | --- | --- |
| 1 | | $14.16 \pm 3.2 \mu\text{M}$ | $9.5 \pm 1.6 \mu\text{M}$ |
| 5 | | $36.47 \pm 4.8 \mu\text{M}$ | NA |
| 10 | | $39.5 \pm 7.1 \mu\text{M}$ | $105.8 \pm 12.65 \mu\text{M}$ |
| 24 | | $2.21 \pm 1.4 \mu\text{M}$ | $22.3 \pm 6.1 \mu\text{M}$ |
| 28 | | $426.93 \pm 29.2 \mu\text{M}$ | $411.3 \pm 51.7 \mu\text{M}$ |
| 50 | | $42.66 \pm 6.7 \mu\text{M}$ | NA |
| 58 | | $85.13 \pm 12.7 \mu\text{M}$ | NA |
| 63 | | $40.98 \pm 4.8 \mu\text{M}$ | NA |
| 64 | | $66.48 \pm 14.7 \mu\text{M}$ | NA |
| 68 | | $4.05 \pm 3.2 \mu\text{M}$ | $1.32 \pm 0.8 \mu\text{M}$ |
| 100 | | $343.78 \pm 22.6 \text{ nM}$ | $231.4 \pm 34 \text{ nM}$ |
| 104 | | $407.08 \pm 14.9 \text{ nM}$ | $840.6 \pm 112.8 \text{ nM}$ |

### Binding Mode Analysis

Docking Analysis. Following the biological evaluation, compounds **100** and **104** emerged as the most active derivatives. Interestingly, these two molecules belong to two distinct clusters among those previously defined based on shape.

Specifically, compound **100** belongs to cluster 3 (Figure 7). In this predicted binding mode, the 6-chloroindolizine moiety is oriented toward the H38 side chain (Figure 7a), allowing for the potential formation of halogen bond interactions. The spatial arrangement of this aromatic portion also might allow the formation of favorable cation-π interactions with the K95 side chain. Moreover, the carbonyl group adjacent to the pyrrolidine establishes H-bond interactions with the side chains of K109 and K2. This latter residue side chain might also potentially engage in cation-π interactions with the central triazole ring. Finally, the terminal carboxylic acid forms H-bond interactions with the D106 backbone N and a coulombic contact with K2.

**Figure 7.**
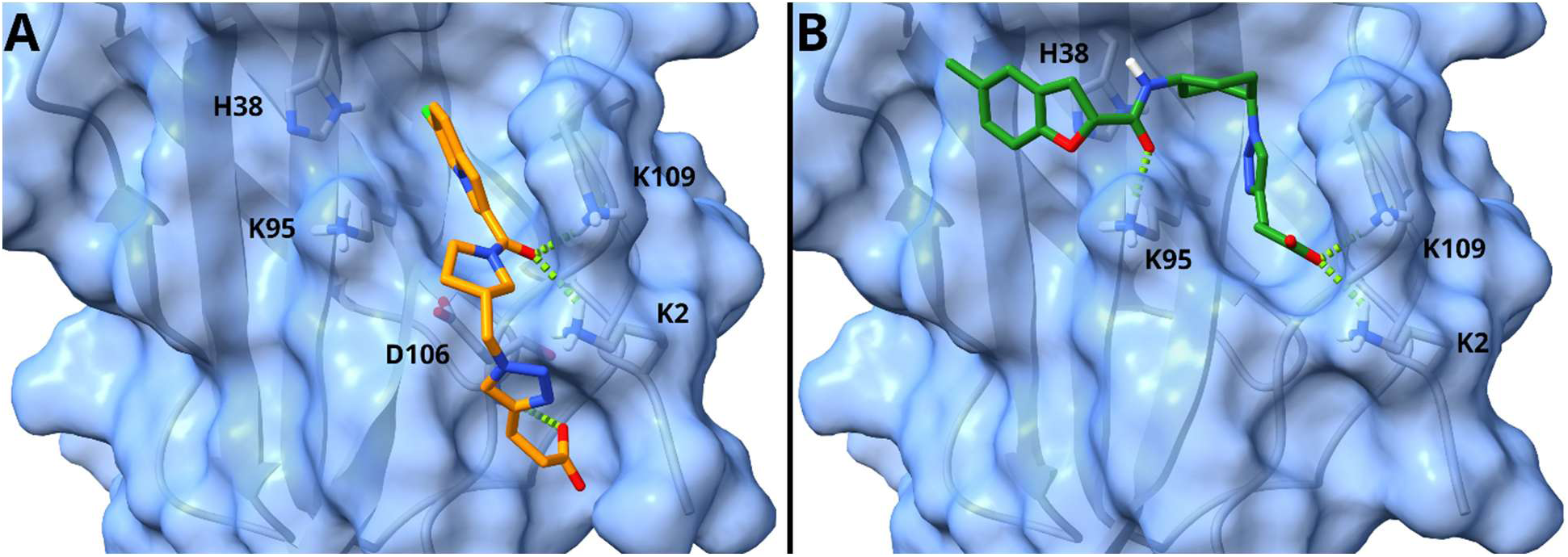
Predicted binding mode of both **100** (a) and **104** (b), orange and green sticks respectively, in complex with the X-Ray structure of CD28. The protein is depicted as grey ribbons and sticks and a blue surface.

Conversely, compound **104** was assigned to cluster 1. In its predicted binding pose, the 5-methylbenzofuran group forms a T-shaped interaction with the H38 side chain, while the cyclohexylamide CO H-bonds with the K95 side chain. This K95 residue may also mediate cation-π interactions with the triazole core of **104**. Similar to **100**, the terminal carboxylic acid of **104** is involved in strong ionic interactions with both K109 and K2 side chains.

All in all, given the high structural similarity between the two molecules, their convergent interaction profile with the same set of amino acid residues within the targeted pocket, and their comparable biological activities, it is tempting to postulate that both molecules might be able to alternatively adopt either binding mode. Therefore, for future hit-to-lead optimization campaigns, both binding modes (represented by clusters 1 and 3 in Figure 3) might be considered as highly valid and acceptable conformations for the design of novel CD28 binders.

### Cell-Based Reporter Assay Confirms CD28 Signaling Blockade by Lead Compounds

To assess whether biochemical disruption of CD28–CD80 binding translated into suppression of CD28-dependent signaling in cells, ELISA-active compounds were evaluated using a CD28 Blockade Bioassay. This luciferase-based reporter assay employs Jurkat effector cells co-cultured with Raji artificial antigen-presenting cells (aAPCs) and quantitatively measures CD28-mediated costimulatory signaling. Compounds were tested in a dose–response format, and luminescence was recorded after 5 h of incubation. IC₅₀ values were derived from three independent experiments.

Of the seven compounds that showed dose-dependent inhibition in the CD28–CD80 ELISA assay, three demonstrated reproducible, concentration-dependent suppression of CD28-driven reporter activity. **100** exhibited the strongest cellular potency (IC₅₀ = 748.89 ± 124.7 nM), followed by 104 (IC₅₀ = 1.13 ± 0.21 μM) (Figure 8A-B). **24** showed weaker but measurable activity (IC₅₀ = 60.7 ± 11.1 μM) (Figure 8C). The remaining ELISA-active compounds did not exhibit dose-dependent inhibition in this cellular assay, indicating that biochemical competition does not consistently translate into effective blockade of CD28-mediated signaling. Across the screening cascade, **100** and **104** consistently demonstrated high-affinity binding (MST), potent disruption of CD28-CD80 interaction (ELISA), and submicromolar inhibition of CD28-driven reporter signaling, establishing them as the most advanced CD28 antagonists within the series.

**Figure 8.**
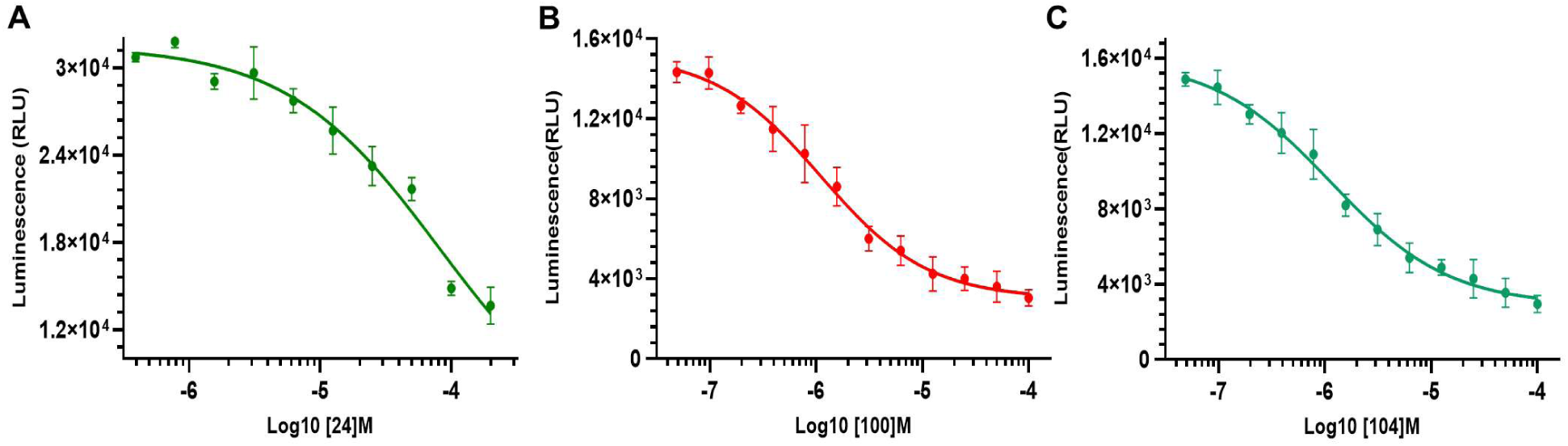
Dose–response curves from a CD28 Blockade Bioassay performed using Jurkat effector cells co-cultured with Raji antigen-presenting cells. Luminescence (RLU) was measured following compound treatment over a defined concentration range using serial dilutions. Data are presented as mean ± SEM. Solid lines represent nonlinear regression fits used to determine IC₅₀ values. (A) **24**, (B) **100**, and (C) **104**.

### Assessment of T Cell Activation in a Tumor-PBMC 3D Co-culture Model

To evaluate the functional impact of our lead CD28 antagonists (**100** and **104**) under physiologically relevant conditions, we employed a 3D co-culture system consisting of IFN-γ-conditioned A549 tumor spheroids combined with primary human PBMCs. T cell receptor signaling was initiated using submaximal anti-CD3 stimulation to ensure that downstream activation remained dependent on CD28-mediated co-stimulatory signals provided by tumor-associated CD80/CD86.

This setup produced strong T cell activation (Figure 9a,b), as demonstrated by increased release of hallmark cytokines including IFN-γ, IL-2, and soluble CD69. Both **100** and **104** attenuated cytokine production in a dose-responsive manner (1, 5, and 10 μM), confirming effective interference with CD28-driven co-stimulation. Across all measured endpoints, **100** showed slightly greater inhibitory potency than **104**, in agreement with its superior biochemical activity profile. The clinical-stage CD28 antagonist FR104 was included as a benchmark control. Collectively, these findings demonstrate that both compounds suppress CD28-dependent T cell activation within a human tumor-immune microenvironment model, with **100** representing the more advanced candidate for further development.

**Figure 9.**
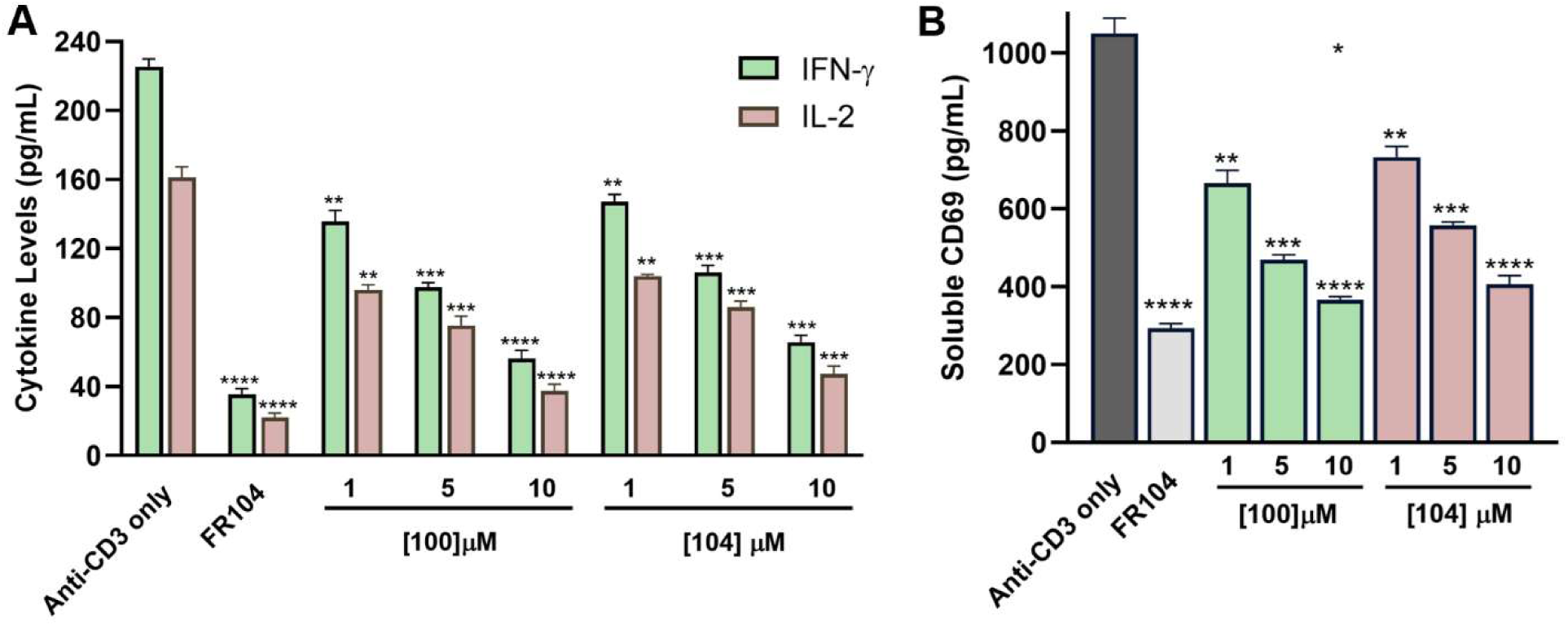
Concentration-dependent suppression of T cell activation in a tumor-PBMC co-culture model. **(A)** IFN-γ and IL-2 concentrations were measured in supernatants collected after 48 h of co-culture between A549 tumor spheroids and human PBMCs (effector-to-target ratio 5:1). Cultures were stimulated with anti-CD3 (0.3 µg/mL) and treated with FR104 (10 µg/mL), **100** (1, 5, or 10 µM), or **104** (1, 5, or 10 µM). **(B)** Soluble CD69 levels were determined by ELISA under identical experimental conditions following 48 h of co-culture. Data are presented as mean ± SD from three independent wells. Statistical analysis was conducted relative to the anti-CD3-stimulated control group using one-way ANOVA with Dunnett’s multiple comparison test. Significance levels are indicated as follows: ** *p* < 0.01; *** *p* < 0.001; **** *p* < 0.0001 versus anti-CD3 alone.

### Human PBMC-Airway Tissue Co-culture Model

To further examine the biological activity of **100** (the most potent lead from this study) in a physiologically relevant setting, we implemented a human co-culture platform designed to mimic immune-epithelial interactions at mucosal surfaces. In this system, differentiated airway epithelial tissues (MucilAir™, Epithelix) were combined with primary human PBMCs and activated using plate-bound anti-CD3 together with soluble anti-CD28 to induce co-stimulatory signaling. This configuration preserves important characteristics of the mucosal immune niche, including epithelial barrier function, localized cytokine signaling, and bidirectional communication between immune and epithelial compartments. Cytokines were measured from the apical supernatant after 48 h to determine the impact of CD28 inhibition.

Dual CD3/CD28 stimulation triggered strong production of IFN-γ, IL-2, and TNF-α, reflecting robust T cell activation. Administration of **100** (1, 5, and 10 μM) attenuated cytokine release in a concentration-dependent manner across the three tested doses (Figures 10A-C). FR104 was included as a reference inhibitor to confirm assay responsiveness. Collectively, these data indicate that **100** retains potent inhibitory activity in a primary human mucosal model, consistent with its performance in tumor-PBMC co-cultures, supporting effective CD28 blockade across diverse immune microenvironments.

**Figure 10.**
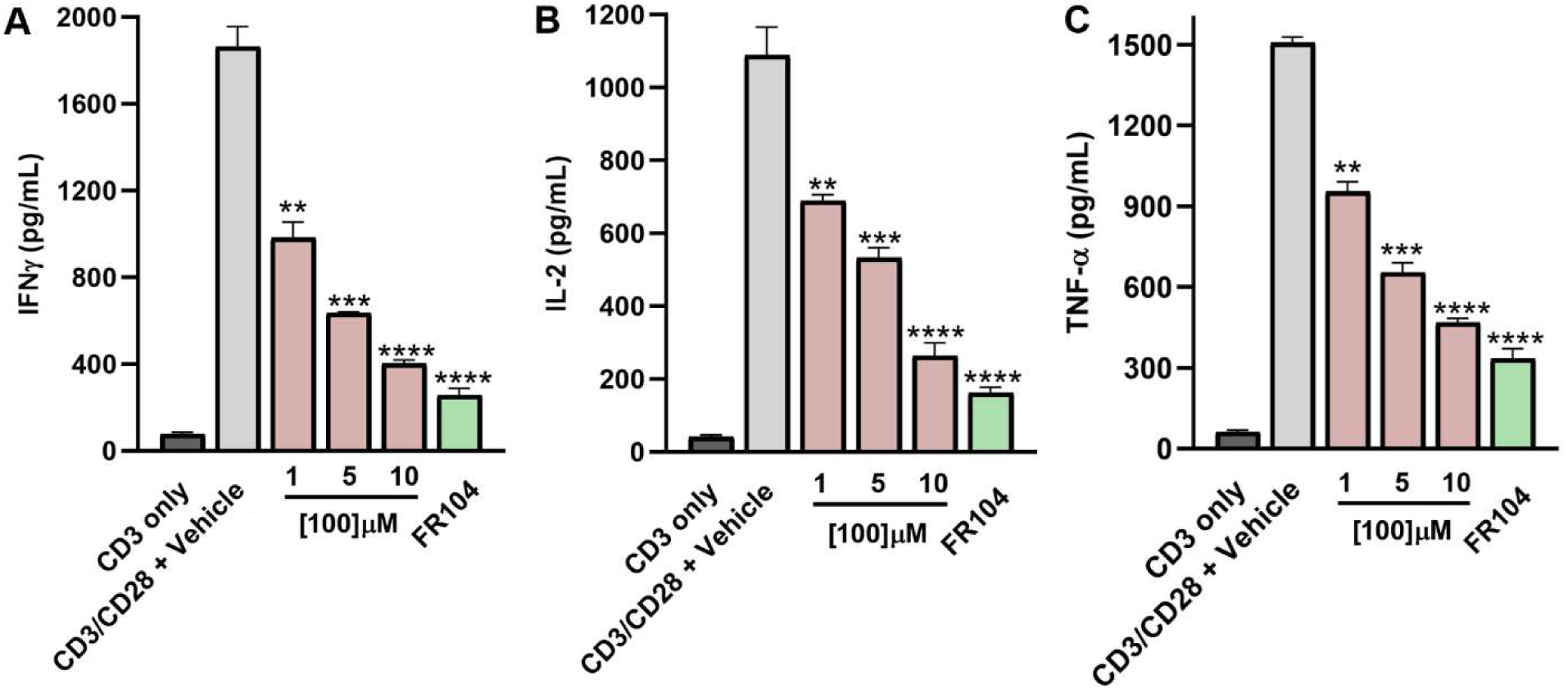
Inhibition of CD28-driven cytokine production by 100 in a human PBMC-airway tissue co-culture system. Primary human PBMCs were applied to differentiated MucilAir™ airway epithelial inserts (Epithelix) and stimulated for 48 h with plate-immobilized anti-CD3 together with soluble anti-CD28. Treatments included vehicle control (0.1% DMSO), **100** at 1, 5, or 10 µM, or the reference CD28 antagonist FR104 (10 µg/mL). Levels of IFN-γ **(A)**, IL-2 **(B)**, and TNF-α **(C)** in apical supernatants were measured by ELISA and expressed in pg/mL. **100** reduced cytokine secretion in a concentration-dependent fashion, with maximal inhibition at 10 µM, approaching the effect observed with FR104. Statistical analysis was performed relative to the CD3/CD28-stimulated vehicle control using one-way ANOVA followed by Dunnett’s multiple comparison test. Significance thresholds are indicated as ** *p* < 0.01, *** *p* < 0.001, and **** *p* < 0.0001 versus CD3/CD28 + vehicle.

## Conclusions

In this study, we successfully addressed the challenge of targeting the shallow, solvent-exposed protein-protein interaction interface of CD28 using small molecules. By taking advantage of the PyRMD2Dock suite developed by some of us, we coupled the screening capabilities of an ML method with the predictive power of docking-based VS, enabling the rapid and efficient virtual screening of 48 million compounds from the Enamine REAL Diversity Set. This represents the first real-world, prospective application of this AI-enforced pipeline. Employing a tailored filtering strategy set of 150 candidates was prioritized for experimental evaluation. As a result, we identified multiple direct CD28 binders, notably compounds **100** and **104**, which exhibited submicromolar affinities (K_d_ = 343.78 nM and 407.08 nM, respectively). These findings highlight the critical advantage of mining ultra-large chemical spaces to identify high-affinity ligands.

These hits demonstrated potent competitive disruption of the CD28-CD80 interaction and efficient blockade of the CD28-mediated signaling in cellular reporter assays. Crucially, the biological efficacy of these compounds translated into complex, physiologically relevant 3D co-culture systems. Both compounds significantly attenuated T cell activation and cytokine secretion in a tumor-PBMC model, while **100** also showed potent inhibitory activity in a mucosal epithelial-PBMC platform. Furthermore, computational analysis revealed that the highly active compounds **100** and **104** should occupy the target cleft in distinct, equally valid binding orientations, offering valuable insights for future hit-to-lead optimization campaigns. Ultimately, the discovery of these potent CD28 antagonists robustly validates the PyRMD2Dock platform. This study demonstrates that AI-driven, large-scale virtual screening can successfully unlock structurally demanding immune receptor interfaces, providing a scalable and highly effective paradigm for targeting complex PPIs

## Material and Methods

### Molecular docking

X-Ray structure of CD28 in complex in complex with the Fab fragment of a mitogenic antibody (1YJD)^6^ was sourced from the RCSB PDB database and underwent preliminary adjustments for docking purposes using the protein preparation wizard integrated into the Schrödinger suite.^18^ The hydrogen atoms were added and minimized, the solvent molecules were removed, and the appropriate protonation and tautomeric state of the protein’s side chains were calculated at physiological pH. Using the AutoDockTools Python scripts, the CD28 structure was converted into the AutoDock PDBQT format, where, compared to a standard PDB file, Gasteiger charges are added to the atoms, and the torsional freedoms of the various bonds are described ^19^. Then, the receptor grid maps were calculated with the AutoGrid4 software, mapping the receptor interaction energies using every AutoDock atom type as a probe, and the docking grid box was centred on the binding site made by H38, F93, K95, D106, N107, K109, and S110 residues (XYZ size of the box: 60 x 60 x 60 with a spacing of 0.375 Å). From the ERDS, a set of 2.4M compounds was selected, and the molecules’ SMILES strings were converted into 3D structures with the employment of the LigPrep routine in Schrödinger’s Maestro suite.^20^ After, their tautomeric and protonation states were calculated at physiological pH, and the possible enantiomers were generated. The prepared molecules were exported as PDB files and converted into PDBQTs by making use of the AutoDock suite scripts. For the docking calculations, attained through AD4-GPU, the Lamarckian Genetic Algorithm (LGA) was employed, encompassing a total of 50 LGA runs.^14^ All other settings were maintained at their default values. The docking results were subsequently grouped based on the RMSD criterion, whereby solutions differing by less than 2.0 Å were considered part of the same cluster. The ranking of these clusters was determined based on the calculated free energy of binding (ΔG_AD4_).

### AI-enforced Virtual Screening

To create a machine learning (ML) prediction model, PyRMD was fed with a comma-separated file (.csv) generated by extracting the predicted lowest ΔG_AD4_ for each of the 2.4 million docked compounds extracted from the ERDS ultra-large database.^13^ This led to the construction and preparation of the training dataset. According to their predicted ΔG_AD4_, the compounds included in the .csv file were classified into three groups: “actives”, “inactives”, and discarded. Compounds whose predicted ΔG_AD4_ falls below the “activity” thresholds of −8.25, −8.0, −7.75, and −7.50 kcal/mol were placed in the active group. Instead, those with a ΔG_AD4_ value higher than the “inactivity” threshold of −4.75, −4.50, −4.25, and −4.0 kcal/mol went into the “inactives” group. Moreover, compounds whose ΔG_AD4_ value falls above the “activity” threshold and “below-the “inactivity” threshold were discarded. By selecting the MinHash fingerprints (MHFP) with a vector length of 2048 units for the featurization process, and by varying the ε cutoff for “actives” (0.70-0.99 with a 0.05 step) and “inactives” (0.60-0.95 with a 0.05 step), 896 different models were generated.^21^ During model training and validation, PyRMD Studio automatically applied Butina clustering coupled with a stratified approach to separate structural clusters between training and test sets. This rigorous data splitting strategy prevented data leakage and ensured a realistic evaluation of the models’ true generalization capabilities

Across all models, PyRMD returns relevant metrics to evaluate their predictive performance (i.e., TPR, FPR, F1-Score, ROC AUC, BED ROC, PRC AUC). In this work, the selected model was chosen by maximizing the True positive rate (TPR)/False positive rate (FPR) trade-off alongside robust Precision and F-score metrics. Once the model was generated, PyRMD was used to screen the remaining ∼46 million compounds from the ERDS database and it automatically returned all the compounds deemed to be active along with a confidence score of its prediction (RMD Score).

### Compound selection and procurement

Following the computational pipeline described previously (see Results and Discussion), a total of 150 compounds from the ERDS ultra-large database were selected for experimental evaluation. From this set, all the compounds were synthesized on demand by Enamine (Kyiv, Ukraine) from the REAL Space collection. All compounds were supplied with reported purity ≥90% (LC–MS verified by the supplier) and were used without further purification. Structural identities were confirmed based on supplier-provided analytical data (HPLC and MS). HPLC spectra and ^1H NMR characterization data for all compounds are provided in the Supplementary Information (pp. S2–S11). SMILES, purity information, and vendor codes for all compounds used in this study are listed in the accompanying Sl Excel file.

### Screening of small-molecule binders to human CD28 using the Dianthus TRIC assay

His-tagged recombinant human CD28 extracellular domain (ECD) was obtained commercially and labeled using RED-tris-NTA 2nd Generation dye (NanoTemper Technologies) according to the manufacturer’s instructions. Briefly, CD28 protein was incubated with dye at a 1:0.5 molar ratio in assay buffer (PBS, pH 7.4, supplemented with 0.05% Tween-20) for 30 min at room temperature in the dark.

For primary screening, labeled CD28 was mixed 1:1 with compounds to yield a final compound concentration of 100 μM in 2% DMSO ^22^. Samples were incubated for 15 min at room temperature prior to measurement. 2% DMSO served as the negative control, and BPU11 was included as a reference compound. After brief centrifugation (1000 × g, 1 min), 20 μL of each sample was loaded into premium capillaries and analyzed using the Dianthus NT.23 Pico instrument (NanoTemper Technologies) at 25 °C.

Fluorescence at 670 nm was recorded for 5 s before and 5 s after IR-laser activation. Normalized TRIC signals (Fnorm) were calculated as the ratio Fhot/Fcold. Each compound was tested in triplicate technical replicates, and control samples were distributed throughout the 384-well plate to monitor signal stability.

### Spectral shift–based binding affinity measurements

Binding affinities were determined using a Monolith X instrument (NanoTemper Technologies) operated in spectral shift detection mode. His-tagged recombinant human CD28 extracellular domain (Acro Biosystems) was labeled with RED-tris-NTA 2nd Generation dye according to the manufacturer’s instructions. Labeling was performed in assay buffer (PBS, pH 7.4, containing 0.05% Tween-20) and incubated for 30 min at room temperature in the dark.

Compounds were prepared as serial dilutions spanning concentrations from 500 μM to 1 nM in assay buffer containing 2% DMSO. Labeled CD28 was mixed 1:1 with compound solutions and incubated for 15 min at room temperature in the dark before measurement.

Samples were loaded into Monolith premium capillaries and measured at 25 °C. Binding was quantified by monitoring ligand-induced changes in the fluorescence emission ratio (F670/F650) upon IR-laser excitation. Normalized fluorescence ratio values were plotted against compound concentration to generate binding curves.

Each compound was tested in technical replicates, and binding experiments were independently repeated. Dissociation constants (K_d_) were determined by nonlinear regression using MO. Affinity Analysis software and GraphPad Prism (four-parameter logistic model).

### Evaluation of CD28–CD80 binding inhibition by competitive ELISA

Competitive ELISA assays were performed using a CD28–CD80 binding assay kit (BPS Bioscience) according to the manufacturer’s instructions with minor modifications. Briefly, 96-well plates pre-coated with recombinant human CD28 were equilibrated to room temperature and washed with 1× wash buffer before use.^8, 21^

Serial dilutions of test compounds were prepared in assay buffer containing 2% DMSO. Compounds were co-incubated with biotinylated human CD80 for 1 h at room temperature to allow competitive binding to immobilized CD28. Wells treated with vehicle (2% DMSO) served as negative controls, and wells containing inhibitor buffer without compound were used to define maximal binding.

Following incubation, wells were washed and incubated with streptavidin–HRP diluted in blocking buffer for 1 h at room temperature. After washing, chemiluminescent substrate was added, and luminescence was immediately measured using a microplate reader.

Percent inhibition was calculated relative to vehicle control wells. IC₅₀ values were determined by fitting concentration–response curves using a four-parameter logistic regression model in GraphPad Prism. All experiments were performed in replicates.

### CD28 blockade bioassay (luciferase reporter assay)

Functional inhibition of CD28-mediated signaling was assessed using the CD28 Blockade Bioassay (Promega, Cat. #JA6101) according to the manufacturer’s instructions. Jurkat CD28 Effector Cells (2 × 10⁴ cells per well) were seeded in white 96-well plates and pre-incubated with serial dilutions of test compounds prepared in assay medium containing 1% DMSO ^7^.

Compounds were tested in a concentration–response format using serial dilutions spanning the indicated concentration range. After compound pre-incubation (5 min at 37 °C), Raji artificial antigen-presenting cells (aAPCs; 2 × 10⁴ cells per well) were added to initiate CD28-dependent costimulatory signaling. Plates were incubated for 5 h at 37 °C in a humidified 5% CO₂ atmosphere.

Following incubation, Bio-Glo™ Luciferase Reagent (Promega) was added according to the manufacturer’s protocol, and luminescence was measured using a microplate luminometer. Dose–response curves were generated using GraphPad Prism (four-parameter logistic regression model), and IC₅₀ values were calculated accordingly. Each compound was tested in technical replicates, and experiments were independently repeated. Data are presented as mean ± SEM.

### Assessment of T Cell Activation in a Tumor–PBMC Co-culture Model

To examine the concentration-dependent effects of CD28 blockade in a translationally relevant human system, we utilized a three-dimensional tumor–PBMC co-culture assay. Tumor spheroids were established by plating A549 cells (from ATCC) into ultra-low attachment 96-well plates and allowing aggregation for 48 h. Spheroids were subsequently primed with recombinant human interferon-γ (50 ng/mL) for 24 h prior to immune cell introduction.

Cryopreserved human peripheral blood mononuclear cells (PBMCs) from STEMCELL Technologies were thawed and added to the spheroids at an effector-to-target ratio of 5:1. T cell receptor stimulation was initiated using soluble anti-CD3 antibody (clone OKT3, 0.3 μg/mL). Test compounds were introduced at the onset of co-culture. FR104 (10 μg/mL), an anti-CD28 Fab′ fragment, was included as a reference inhibitor. Compounds **100** and **104** were tested at 1, 5, and 10 μM, with vehicle and anti-CD3-only conditions serving as controls.

After 48 h, culture supernatants were collected for cytokine quantification. IFN-γ and IL-2 levels were measured using human ELISA kits (BioLegend), and soluble CD69 was determined using a human CD69 ELISA kit (Thermo Fisher Scientific).

### CD28-Mediated Cytokine Production in a Human PBMC–Airway Tissue Co-culture System

To investigate the immunomodulatory effects of **100** within a physiologically relevant mucosal setting, a human airway epithelial tissue model (MucilAir™, Epithelix) was co-cultured with PBMCs. Cryopreserved PBMCs (StemCell Technologies) were thawed and allowed to recover for 4 h in RPMI-1640 supplemented with 10% heat-inactivated human AB serum, 2 mM L-glutamine, and 1% penicillin-streptomycin.

MucilAir™ inserts were preconditioned for 24 h in proprietary maintenance medium at 37 °C under 5% CO_2_. PBMCs were adjusted to a density of 2 × 10^6^ cells/mL, and 200 µL was applied to the apical compartment of each insert in 24-well Transwell plates. T cell activation was induced using plate-immobilized anti-CD3 antibody (1 µg/mL) together with soluble anti-CD28 (1 µg/mL). Treatments included vehicle control (0.1% DMSO), **100** at 1, 5, or 10 µM, or the reference CD28 inhibitor FR104 (10 µg/mL).

After 48 h of incubation, apical supernatants were collected and analyzed for IFN-γ, IL-2, and TNF-α using human ELISA kits (BioLegend). Cytokine levels were calculated from standard curves and reported in pg/mL. All experimental conditions were conducted in triplicate across three independent experiments.

## AUTHOR INFORMATION

### DATA AVAILABILITY

The PyRMD Studio software can be downloaded free of charge at https://github.com/cosconatilab/PyRMD-Studio

### ASSOCIATED CONTENT

The following files are available free of charge.

Chemical structures and SMILES strings of the active compounds, HPLC purity, NMR spectra, MST binding studies, and competitive ELISA (PDF).

SMILES strings, Catalog IDs, and purity for all compounds (XLSX).

## Acknowledgments

This work was supported by the National Institute of Diabetes and Digestive and Kidney Diseases (NIDDK) under grant number R01DK137299−PI M.G.

This work was funded by the AIRC (Associazione Italiana per la Ricerca sul Cancro), IG 2021−ID 25865 project−PI S.C.

## Notes

The authors declare no competing financial interest.

## ABBREVIATIONS

PPI: protein-protein interaction
AI: artificial intelligence
VS: virtual screening
SBVS: structure-based virtual screening
LBVS: ligand-based virtual screening
ML: machine learning
AD4-GPU: AutoDock-GPU
ECFP6: Extended Connectivity Fingerprints with a radius of 3
ERDS: Enamine REAL Diversity Set
SASA: solvent-accessible surface area
ECD: extracellular domain
MST: microscale thermophoresis
aAPCs: artificial antigen-presenting cells
PBMC: peripheral blood mononuclear cells
LGA: Lamarckian Genetic Algorithm
MHFP: MinHash fingerprints
TPR: True positive rate
FPR: False positive rate
TRIC: Temperature-related intensity change

## Table of Contents

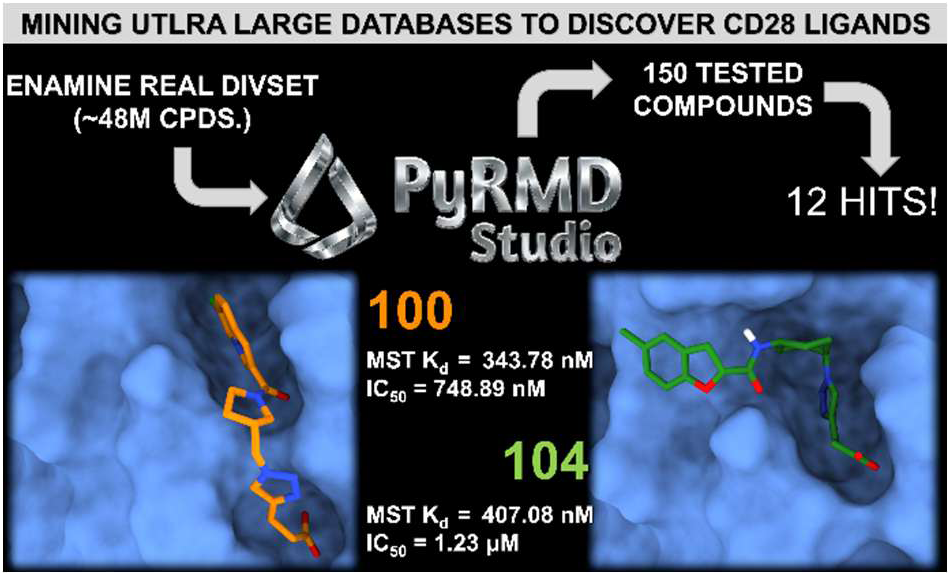

## Notes

### Competing Interest Statement

The authors have declared no competing interest.

