## Supporting Information for "AI-Enforced Ultra-Large Virtual Screening Discovers Potent CD28 Binders"

**Table of Content:**

| **S. No.** | **Content** | **Page Number** |
| --- | --- | --- |
| Table S1 | Structures and vendor codes for all the purchased compounds. | S2-S10 |
| Fig S1-S12 | HPLC Spectra for Active compounds | S10-S14 |
| Fig S13-S18 | 1H-NMR Spectra for Active compounds | S15-S18 |
| Fig S19 | Spectral shift binding curves of **28** and **58** to CD28. | S18 |
| Fig S20 | ELISA dose–response curve of **28** | S19 |

**Table S1.** Structures and vendor codes for all the purchased compounds.

| **Compound#** | **Structure** | **Vendor ID** |
| --- | --- | --- |
| **1** | 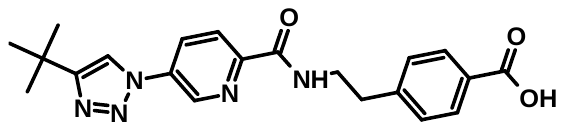 | Z9698696370 |
| **2** | 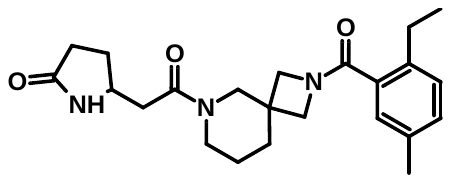 | Z9698696424 |
| **3** | 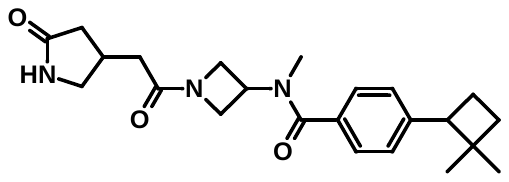 | Z9698696369 |
| **4** | 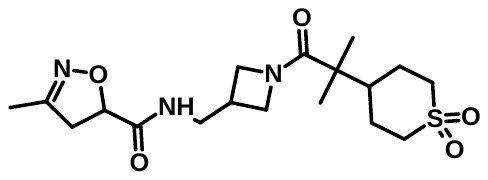 | Z9698696308 |
| **5** | 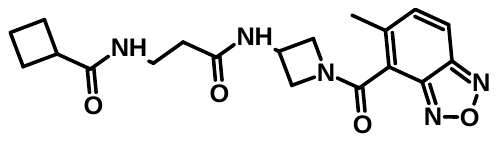 | Z9698696375 |
| **6** | 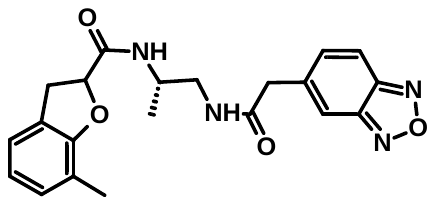 | Z9698696371 |
| **7** | 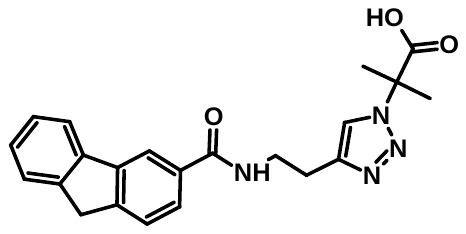 | Z9698696376 |
| **8** | 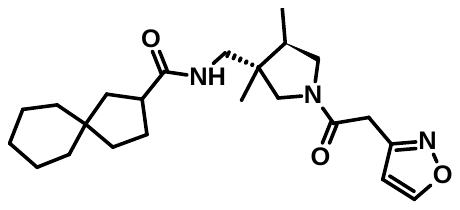 | Z9698696298 |
| **9** | 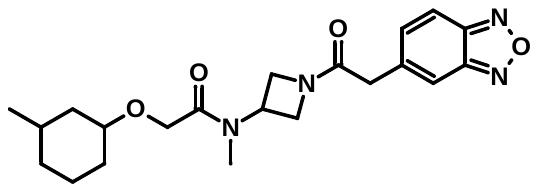 | Z9698696395 |
| **10** | 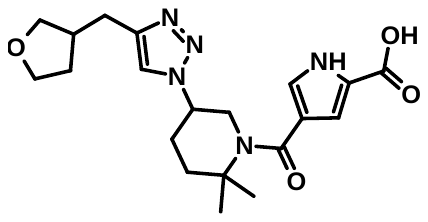 | Z9698696401 |
| **11** | 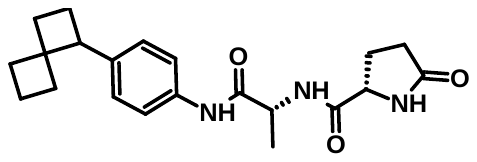 | Z7226927075 |
| **12** | 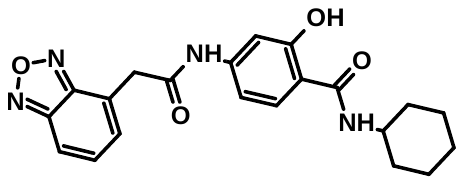 | Z9698696382 |
| **13** | 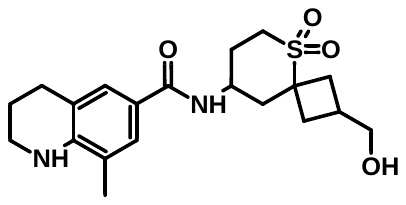 | Z6327159723 |
| **14** | 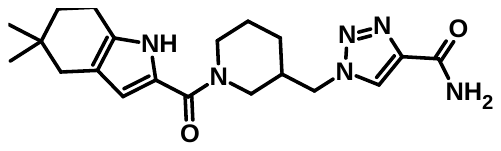 | Z9698696385 |
| **15** | 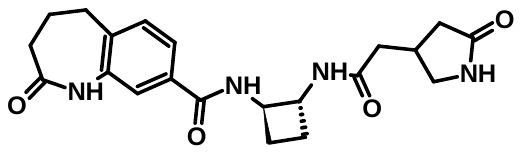 | Z9698696394 |
| **16** | 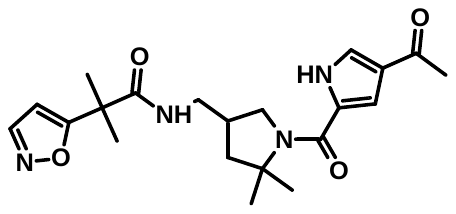 | Z9698696409 |
| **17** | 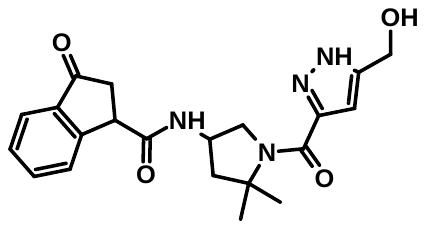 | Z9698696412 |
| **18** | 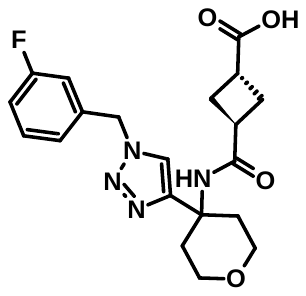 | Z9698696408 |
| **19** | 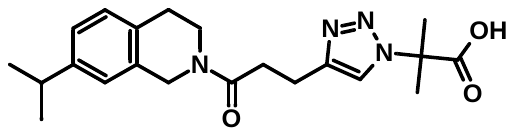 | Z9698696379 |
| **20** | 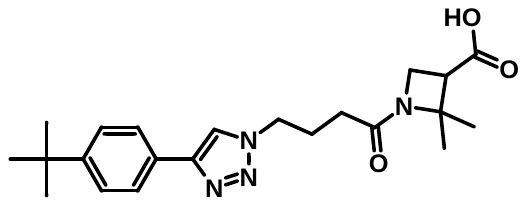 | Z9698696358 |
| **21** | 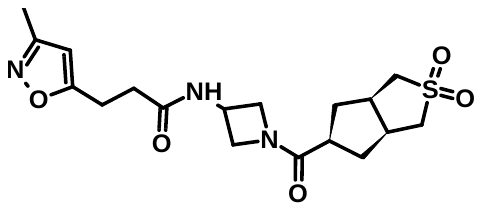 | Z9698696377 |
| **22** | 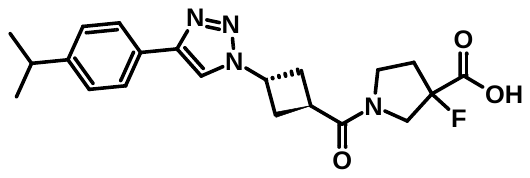 | Z9698696344 |
| **23** | 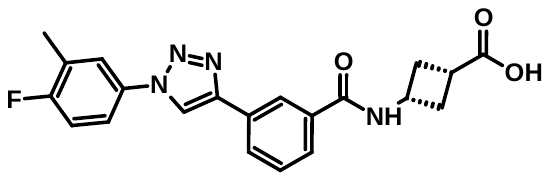 | Z9698696350 |
| **24** | 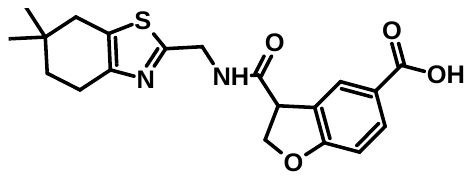 | Z9698696341 |
| **25** | 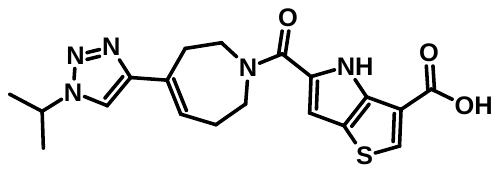 | Z9698696292 |
| **26** | 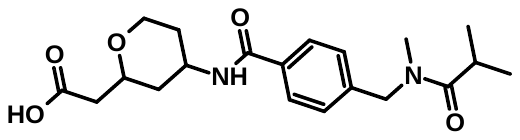 | Z3510250611 |
| **27** | 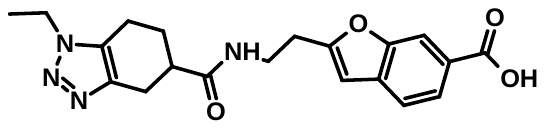 | Z9698696354 |
| **28** | 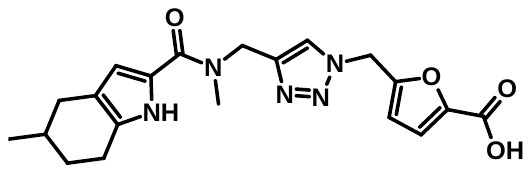 | Z9698696337 |
| **29** | 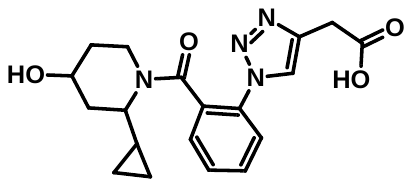 | Z9698696303 |
| **30** | 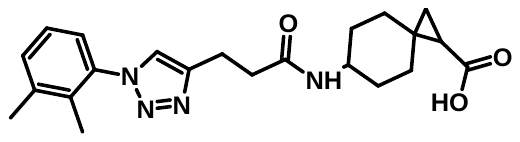 | Z9698696268 |
| **31** | 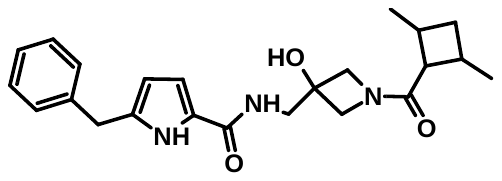 | Z9698696391 |
| **32** | 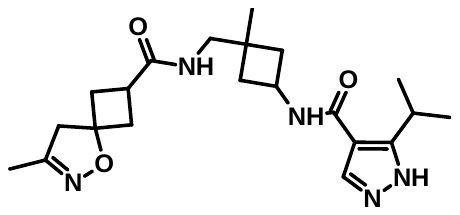 | Z9698696302 |
| **33** | 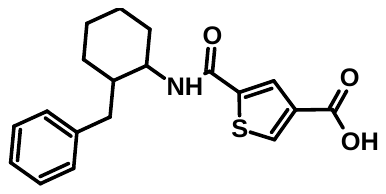 | Z9698696284 |
| **34** | 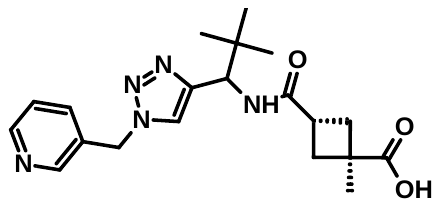 | Z9698696404 |
| **35** | 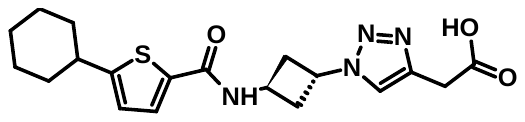 | Z9698696365 |
| **36** | 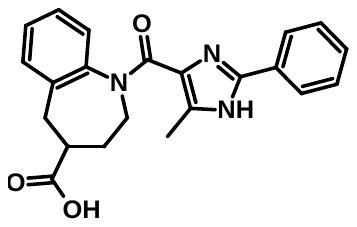 | Z9698696281 |
| **37** | 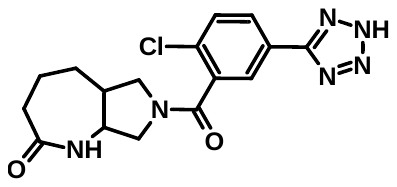 | Z9698696301 |
| **38** | 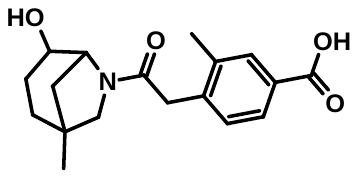 | Z3509247428 |
| **39** | 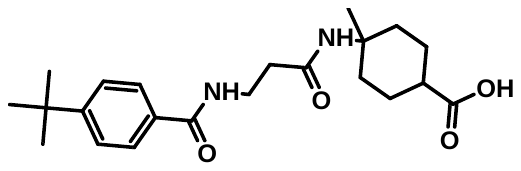 | Z9698696362 |
| **40** | 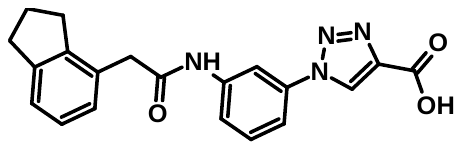 | Z9698696429 |
| **41** | 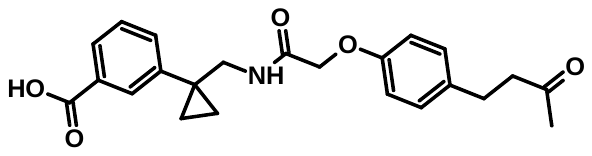 | Z4577965976 |
| **42** | 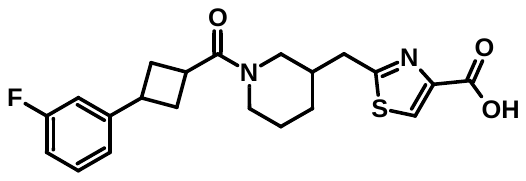 | Z9698696359 |
| **43** | 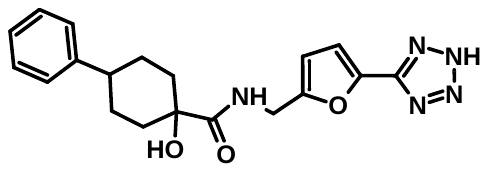 | Z9698696378 |
| **44** | 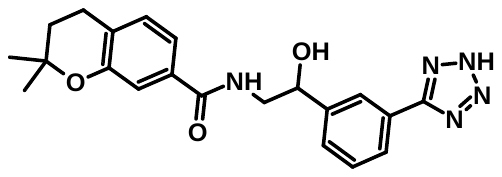 | Z9698696363 |
| **45** | 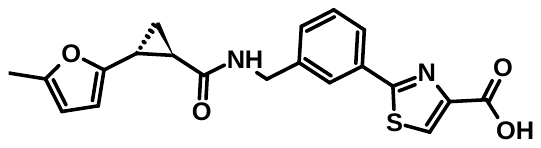 | Z9698696348 |
| **46** | 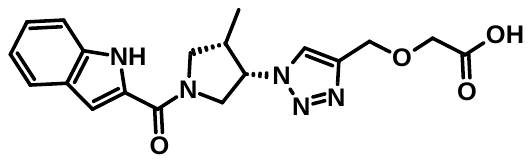 | Z9698696416 |
| **47** | 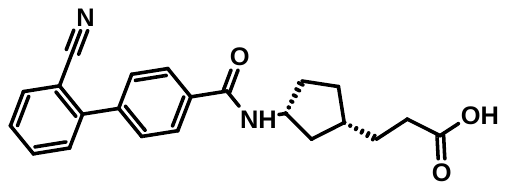 | Z9698696396 |
| **48** | 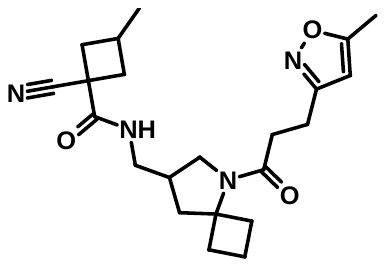 | Z9698696418 |
| **49** | 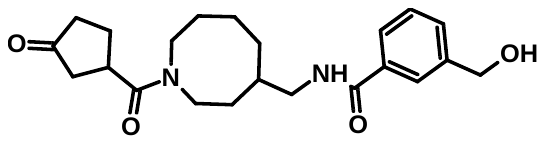 | Z9698696410 |
| **50** | 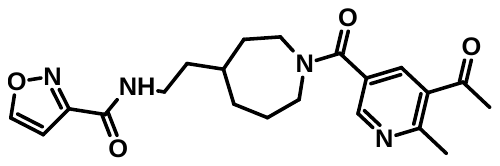 | Z9698696427 |
| **51** | 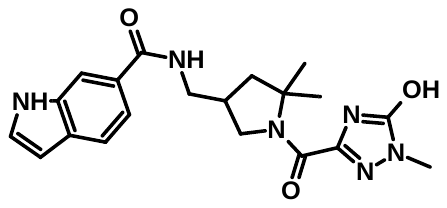 | Z9698696405 |
| **52** | 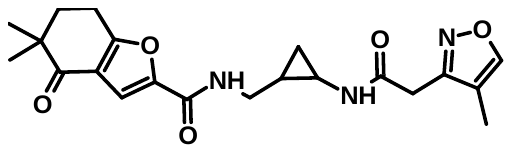 | Z9698696398 |
| **53** | 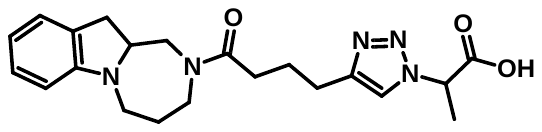 | Z9698696272 |
| **54** | 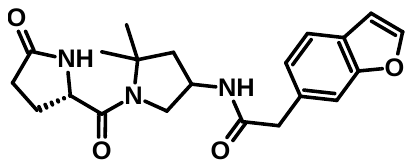 | Z9698696413 |
| **55** | 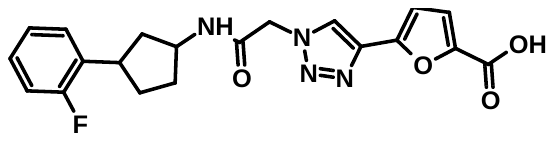 | Z9698696345 |
| **56** | 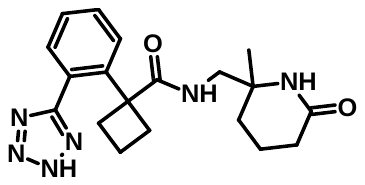 | Z9698696290 |
| **57** | 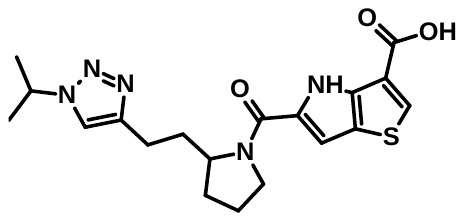 | Z9698696289 |
| **58** | 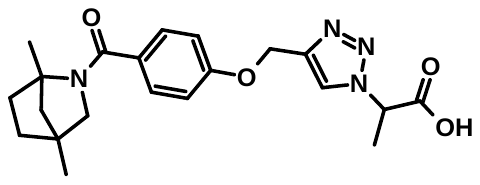 | Z9698696388 |
| **59** | 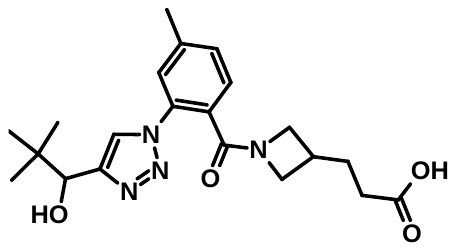 | Z9698696293 |
| **60** | 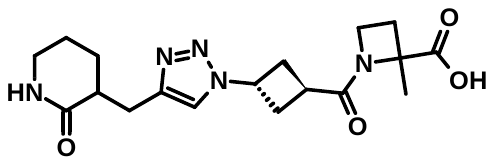 | Z9698696264 |
| **61** | 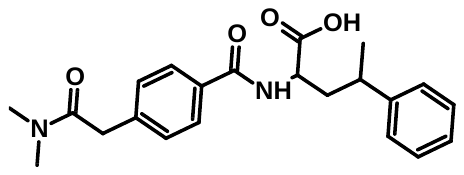 | Z4590802640 |
| **62** | 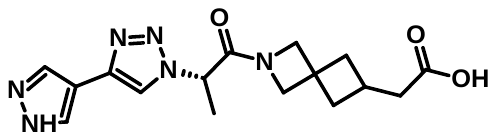 | Z9698696274 |
| **63** | 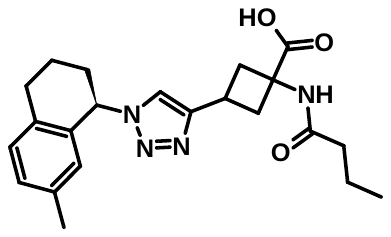 | Z9698696383 |
| **64** | 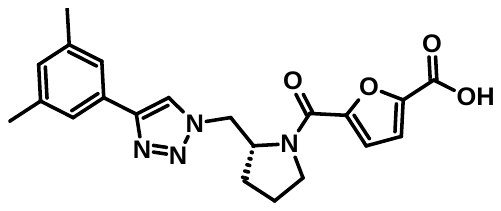 | Z9698696349 |
| **65** | 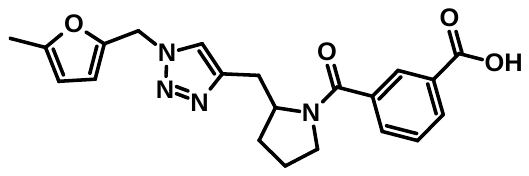 | Z9698696417 |
| **66** | 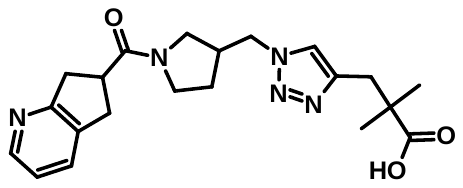 | Z9698696360 |
| **67** | 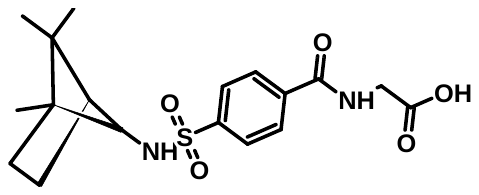 | Z9698696299 |
| **68** | 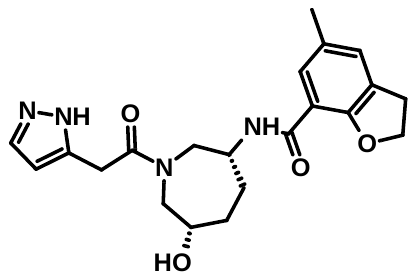 | Z9698696403 |
| **69** | 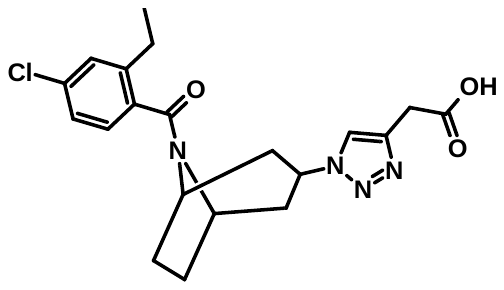 | Z9698696275 |
| **70** | 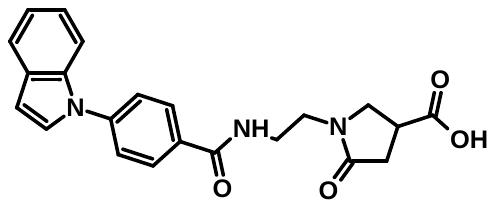 | Z9698696361 |
| **71** | 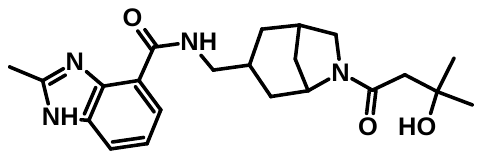 | Z9698696423 |
| **72** | 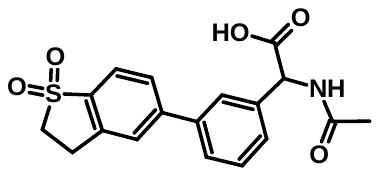 | Z9698696265 |
| **73** | 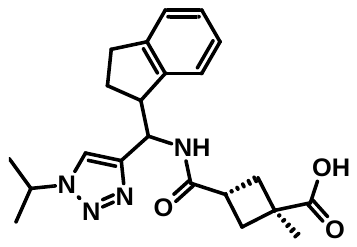 | Z9698696279 |
| **74** | 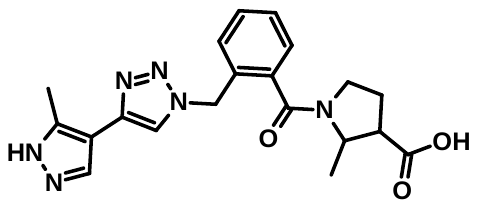 | Z9698696277 |
| **75** | 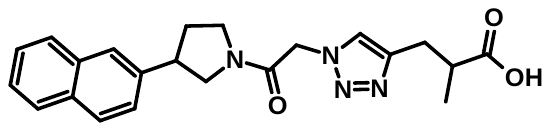 | Z9698696353 |
| **76** | 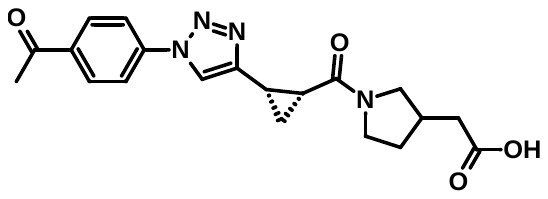 | Z9698696386 |
| **77** | 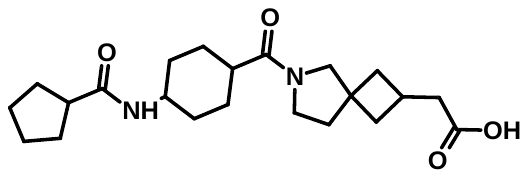 | Z9698696273 |
| **78** | 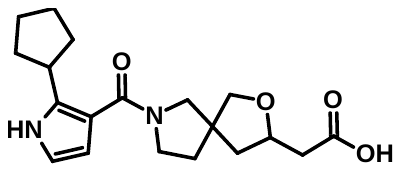 | Z5074828592 |
| **79** | 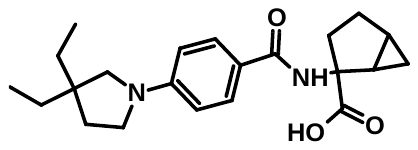 | Z9698696306 |
| **80** | 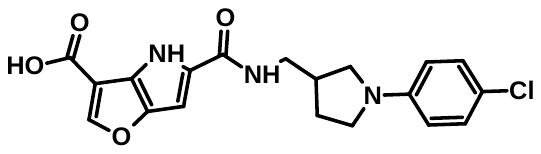 | Z9698696305 |
| **81** | 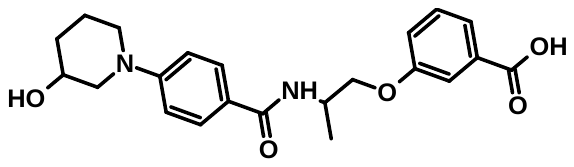 | Z6645588208 |
| **82** | 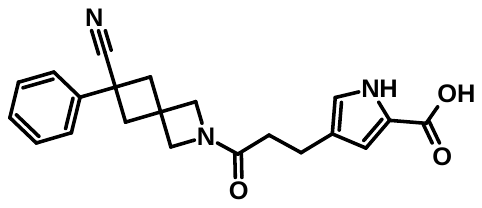 | Z6654658182 |
| **83** | 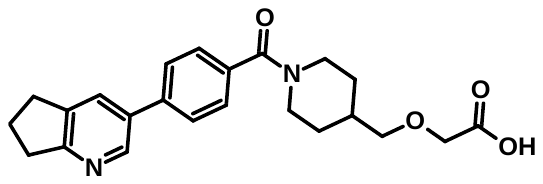 | Z6704781071 |
| **84** | 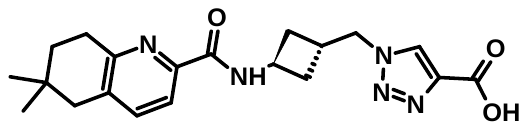 | Z9698696387 |
| **85** | 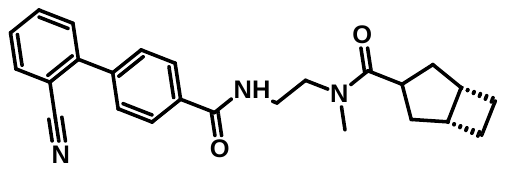 | Z9698696392 |
| **86** | 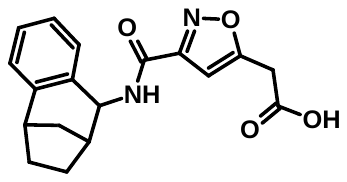 | Z5643051772 |
| **87** | 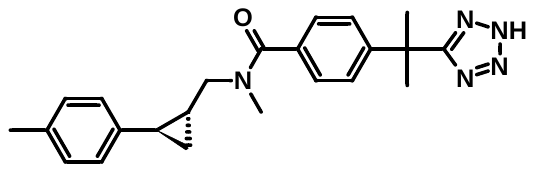 | Z9698696351 |
| **88** | 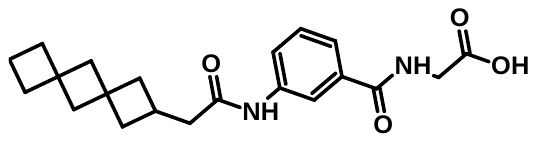 | Z9698696267 |
| **89** | 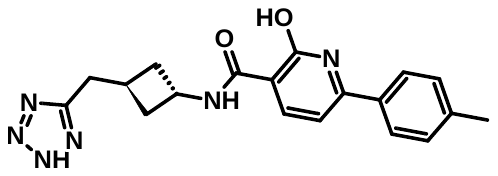 | Z9698696346 |
| **90** | 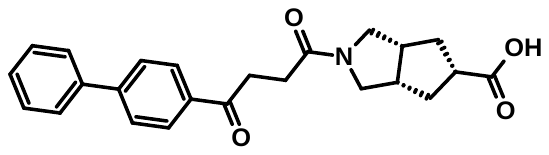 | Z6671929133 |
| **91** | 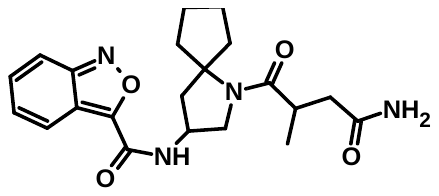 | Z9698696430 |
| **92** | 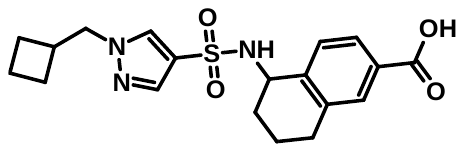 | Z9698696285 |
| **93** | 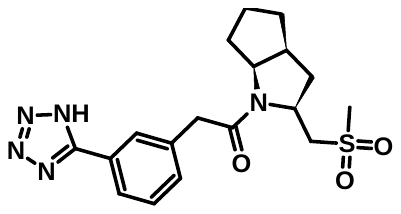 | Z8173342514 |
| **94** | 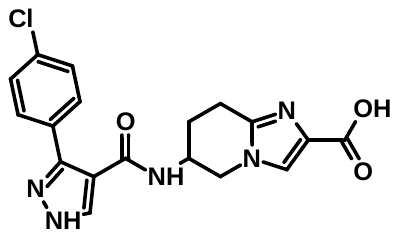 | Z9916238374 |
| **95** | 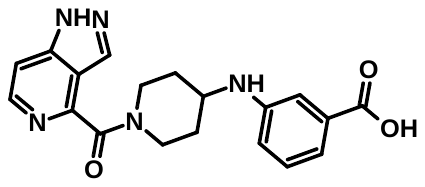 | Z9916238372 |
| **96** | 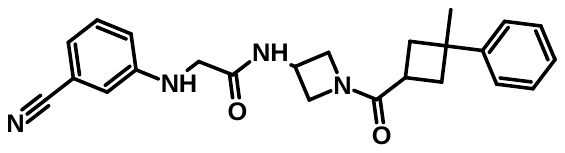 | Z9698696397 |
| **97** | 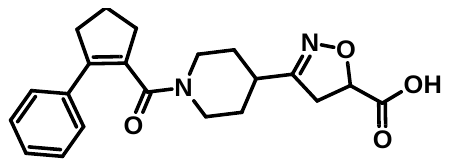 | Z6683469397 |
| **98** | 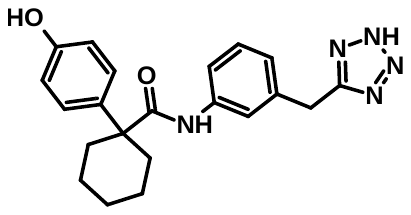 | Z9698696407 |
| **99** | 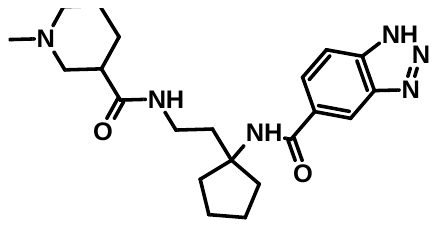 | Z9698696411 |
| **100** | 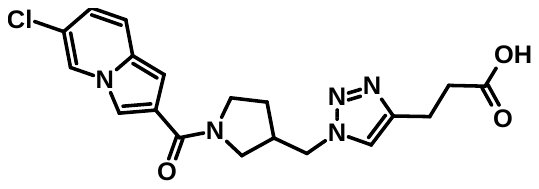 | Z9698696355 |
| **101** |  | Z9698696415 |
| **102** |  | Z9698696374 |
| **103** |  | Z9698696425 |
| **104** |  | Z9698696402 |
| **105** |  | Z9698696426 |
| **106** |  | Z9698696352 |
| **107** |  | Z9698696347 |
| **108** |  | Z9698696283 |
| **109** |  | Z9698696276 |
| **110** |  | Z9698696286 |
| **111** |  | Z9698696357 |
| **112** |  | Z6661443394 |
| **113** |  | Z6706676830 |
| **114** |  | Z9916238373 |
| **115** |  | Z9918750513 |
| **116** |  | Z9698696393 |
| **117** |  | Z9698696307 |
| **118** |  | Z9698696373 |
| **119** |  | Z9698696364 |
| **120** |  | Z9698696421 |
| **121** |  | Z1991870008 |
| **122** |  | Z5273326489 |
| **123** |  | Z6682428204 |
| **124** |  | Z9698696333 |
| **125** |  | Z9698696338 |
| **126** |  | Z9698696255 |
| **127** |  | Z9698696258 |
| **128** |  | Z4577476016 |
| **129** |  | Z6096789177 |
| **130** |  | Z6242298398 |
| **131** |  | Z6681383318 |
| **132** |  | Z9698696253 |
| **133** |  | Z9698696254 |
| **134** |  | Z9698696259 |
| **135** |  | Z9698696260 |
| **136** |  | Z9698696261 |
| **137** |  | Z9698696262 |
| **138** |  | Z9698696321 |
| **139** |  | Z9698696322 |
| **140** |  | Z9698696332 |
| **141** |  | Z9698696334 |
| **142** |  | Z9698696336 |
| **143** |  | Z9698696372 |
| **144** |  | Z9698696339 |
| **145** |  | Z9698696340 |
| **146** |  | Z9698696295 |
| **147** |  | Z9698696343 |
| **148** |  | Z9698696399 |
| **149** |  | Z9698696400 |
| **150** |  | Z5292659966 |

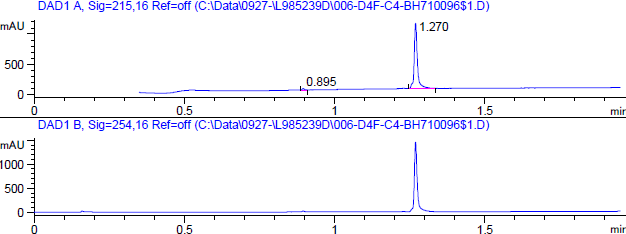

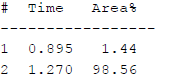

FigS1: HPLC Spectra for **1.**

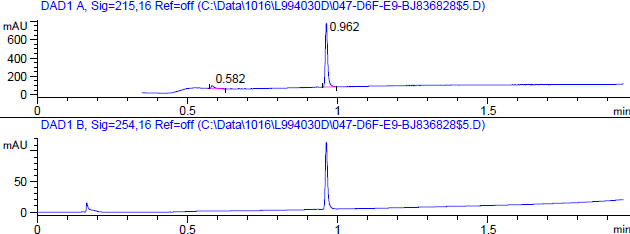

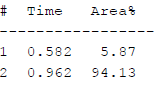

FigS2: HPLC Spectra for **5**

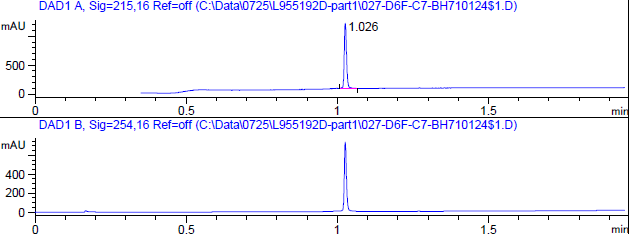

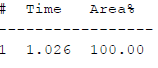

Fig S3: HPLC Spectra for **10**

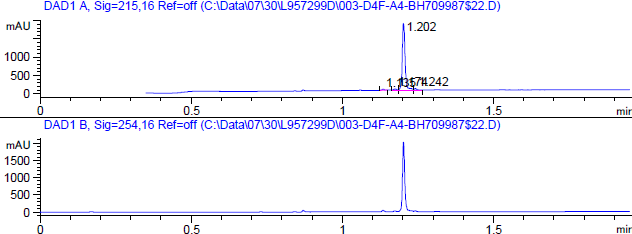

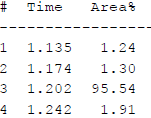

Fig S4: HPLC Spectra for **24**

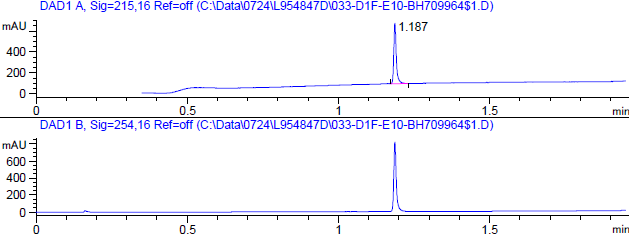

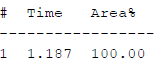

Fig S5: HPLC Spectra for **28**

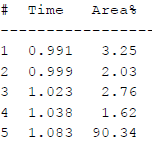

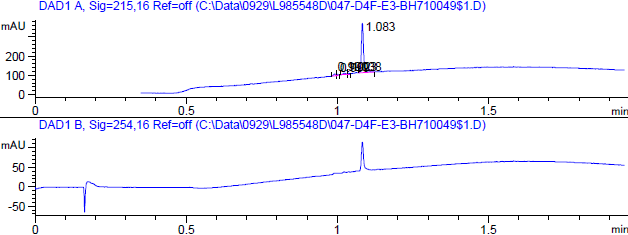

Fig S6: HPLC Spectra for **50**

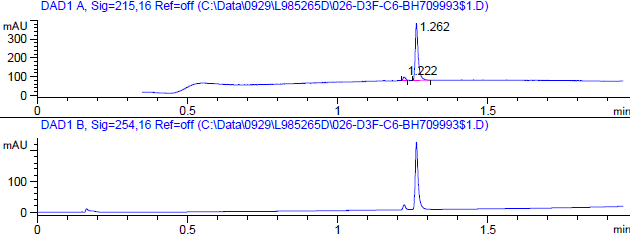

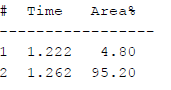

Fig S7: HPLC Spectra for **58**

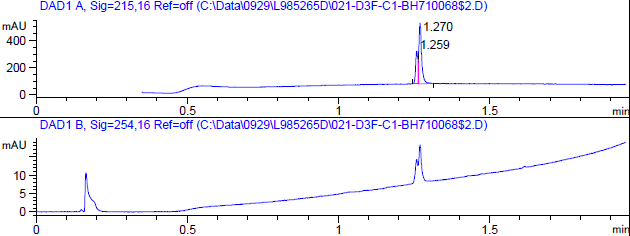

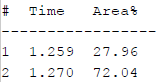

Fig S8: HPLC Spectra for **63**

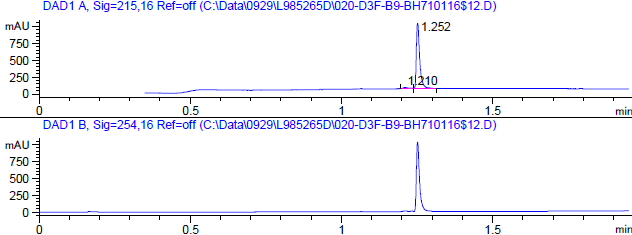

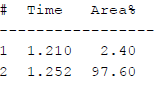

Fig S9: HPLC Spectra for **64**

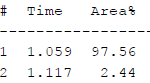

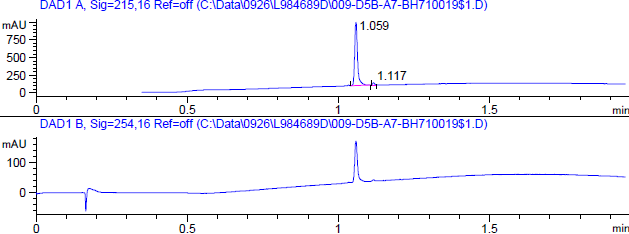

Fig S10: HPLC Spectra for **68**

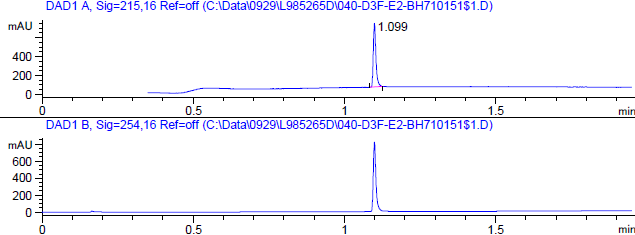

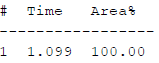

Fig S11: HPLC Spectra for **100**

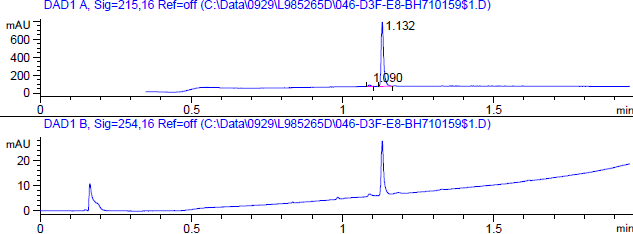

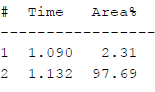

Fig S12: HPLC Spectra for **104**

Fig S13: 1H-NMR Spectra for **24**

Fig S14: 1H-NMR Spectra for **63**

Fig S15: 1H-NMR Spectra for **64**

Fig S16: 1H-NMR Spectra for **68**

Fig S17: 1H-NMR Spectra for **100**

Fig S18: 1H-NMR Spectra for **104**

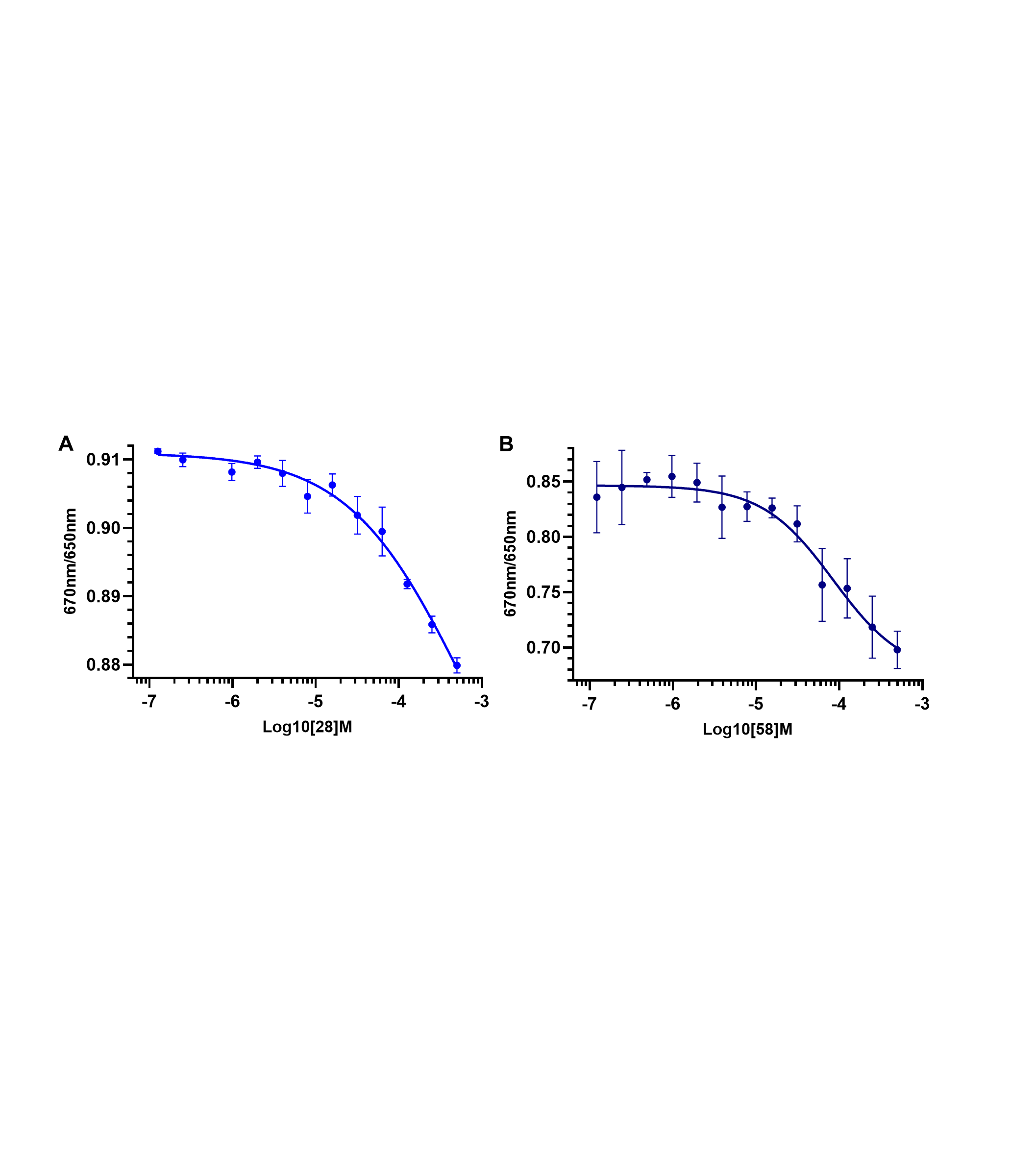

Fig S19: Dose–response curves obtained using Monolith X (F670/F650 spectral shift mode). His-tagged CD28 was incubated with serial dilutions of compounds, and normalized fluorescence ratios were plotted against concentration. Data represent mean ± SEM. Solid lines indicate nonlinear regression fits used to calculate dissociation constants.(A) AU28, Kd = 426.93 μM and (B) AU58, Kd = 85.14 μM.

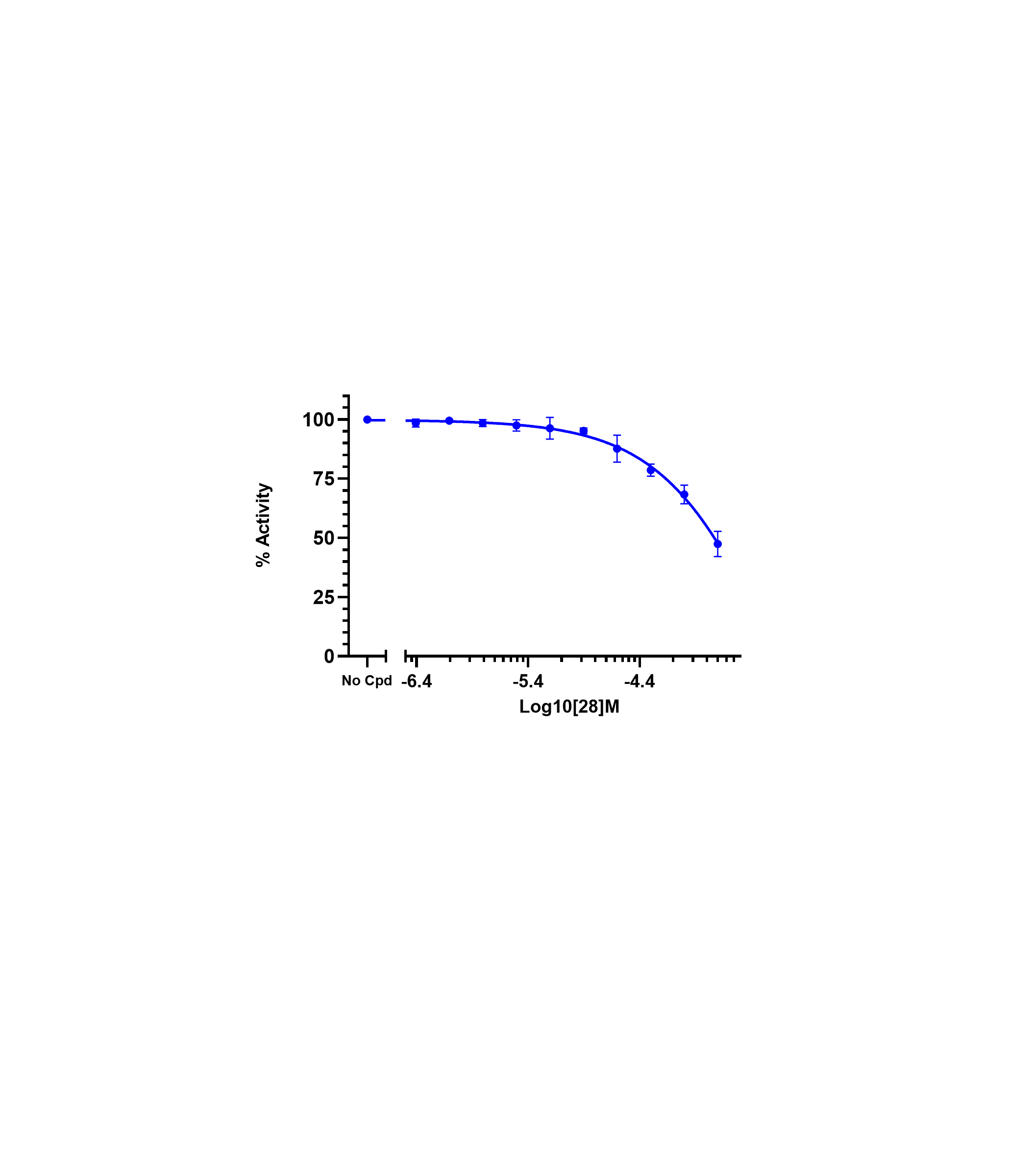

Fig S20. Dose–response inhibition curve of AU28 in a competitive CD28–CD80 ELISA, with an IC₅₀ of 411.3 μM determined by nonlinear regression.
